# Refeeding-associated AMPK_γ1_ complex activity is a hallmark of health and longevity

**DOI:** 10.1101/2022.12.05.519139

**Authors:** Roberto Ripa, Eugen Ballhysa, Joachim D. Steiner, Andrea Annibal, Nadine Hochhard, Christian Latza, Luca Dolfi, Chiara Calabrese, Anna M. Meyer, M. Cristina Polidori, Roman-Ulrich Müller, Adam Antebi

## Abstract

Late-life-initiated dietary interventions negligibly extend longevity or reduce frailty, yet the reason remains unknown. We investigated the age-related changes associated with the fasting response in adipose tissue of the short-lived killifish *N. furzeri*. Transcriptomic analysis revealed the presence of a fasting-like transcriptional program (FLTP) in old animals that is irrespective of their nutritional status and characterized by widespread suppression of anabolic processes. FLTP is associated with reduced expression of the AMPK γ1 regulatory subunit. Accordingly, refeeding positively regulates γ1 expression in young but not in old animals. Fish having sustained AMPK_γ1_ activation had no sign of FLTP in old age and exhibited metabolic health and longevity. In humans, we found that γ1 expression declines with age and is associated with multimorbidity and multidimensional frailty risk. Our study highlights the importance of the refeeding arm in promoting health and longevity and identifies the AMPK_γ1_ complex as a potential target to prevent age-related diseases in humans.

## Introduction

Dietary interventions (DIs) that result in the periodic reduction or removal of caloric intake or specific diet components robustly promote health and longevity^1-4^, but impose a lifelong regimen that is not feasible in humans. To minimize such a burden, the question arises whether DIs benefits can still be induced at later time points. However, DIs initiated at old age in model organisms often fail to extend lifespan^5,6^ or can even lead to frailty^7-10^, indicating these interventions are effective up to mid-life but perhaps become detrimental later, yet the reasons remain unclear. As DIs entail cycles of fasting and refeeding, great attention has been given to unraveling the physiological and molecular responses to nutrient removal and reintroduction. However, whether the physiological response to fasting or refeeding becomes impaired in older animals remains unknown. Here, we employed the short-lived turquoise killifish, which lives up to 6-7 months and shows a functional decline remarkably similar to mammals^11-14^, to study the age-related changes in response to food deprivation. By transcriptomic analysis, we observed that adipose tissue of older animals exhibited a fasting-like transcriptional program (FLTP) irrespective of their nutritional status (fed or fasted), characterized by widespread suppression of anabolic processes. At the gene level, we found that the regulatory AMPK subunit γ1 expression was repressed by fasting and induced by re-feeding. Such regulation was lost during aging resulting in chronic suppression of γ1. By genetic interventions, we discovered that promoting AMPK_γ1_ complex activity maintained the correct response to anabolic stimuli in the adipose tissue of older individuals, thus sustaining regenerative processes late in life and ultimately promoting metabolic health and longer life. Importantly, we also found that γ1 expression declines as a function of age and is prognostic of multimorbidity and multidimensional frailty risk in humans.

## Results

### Aging initiates a fasting-like transcriptional program (FLTP) in killifish visceral adipose tissue

We initially investigated how aging influences the physiologic response to food deprivation in vertebrates using the turquoise killifish. We primarily focused on the adipose tissue as it undergoes substantial remodeling under DIs or during aging^15^. To this end, young (7 weeks) and old (18 weeks) adult male killifish were food deprived for five days, while age-matched control male fish were normally fed twice a day and sacrificed two hours after their last meal, together with fasted fish (figure 1A). As aging can impair feeding behavior or intestinal absorption, we determined blood glucose levels (BGL) to validate that young and old control groups were equally fed and did not have impaired intestinal absorption. BGL was comparable in control groups and significantly lower in the fasted groups (Extended data figure 1A). We then performed transcriptome analysis on the visceral adipose tissue, comparing the fasted and fed conditions of young adult (y-fasted/y-fed) and old (o-fasted/o-fed) wild-type fish. We identified 2352 differentially expressed genes (DEGs) in young fasted relative to young fed fish (FDR<0.05, Extended data figure 1B, supplementary table 1). The majority of these DEGs (86%) were downregulated. KEGG enrichment analysis revealed downregulation of several metabolic pathways, including ribosome, TCA cycle, oxidative phosphorylation, glycolysis, and fatty acid synthesis metabolism, among others (Extended Data Figure 1D-E, supplementary table 2), reflecting a reduction in protein synthesis, metabolic rate, and lipogenesis. Concomitantly, fasting induced upregulation of autophagy and negative regulators of cell proliferation such as *Pax6, Apc2*, and *Bmp4*^*16*^ (Extended Data Figure 1D-1E, supplementary table 1). Thus, in line with mammalian data^17^, the physiological response to food deprivation consists of reducing energy expenditure, protein synthesis, and cellular proliferation in killifish. Notably, gene expression changes in response to food deprivation appeared significantly reduced in old fish. In fact, we could identify only 343 DEGs in old fasted relative to old fed fish (Extended data figure 1C, supplementary table 3). To understand this further, we evaluated the scaled expression of differentially regulated genes (FDR<0.05) under fasting in young adult fish across all groups and observed that the expression profile of such genes in old fully fed fish resembled those of fasted fish (Figure 1B). Accordingly, distance matrix analysis indicated the transcriptome of fully fed old animals clustering together with those of fasted groups (Figure 1C). We further validated the expression of some of the top fasting-responsive genes using an independent cohort of fish by qPCR analysis and observed the same pattern again (Extended data figure 1F). Fasting liberates non-esterified fatty acids (NEFAs) as an alternative energy substrate. Consistent with the RNA-seq analysis, old killifish exhibited elevated levels of plasma NEFAs whether fed or fasted (Figure 1D). Thus, taken together, these data indicate the presence of a fasting-like transcriptional program in the adipose tissue of older animals regardless of their nutritional status.

**Figure 1:**
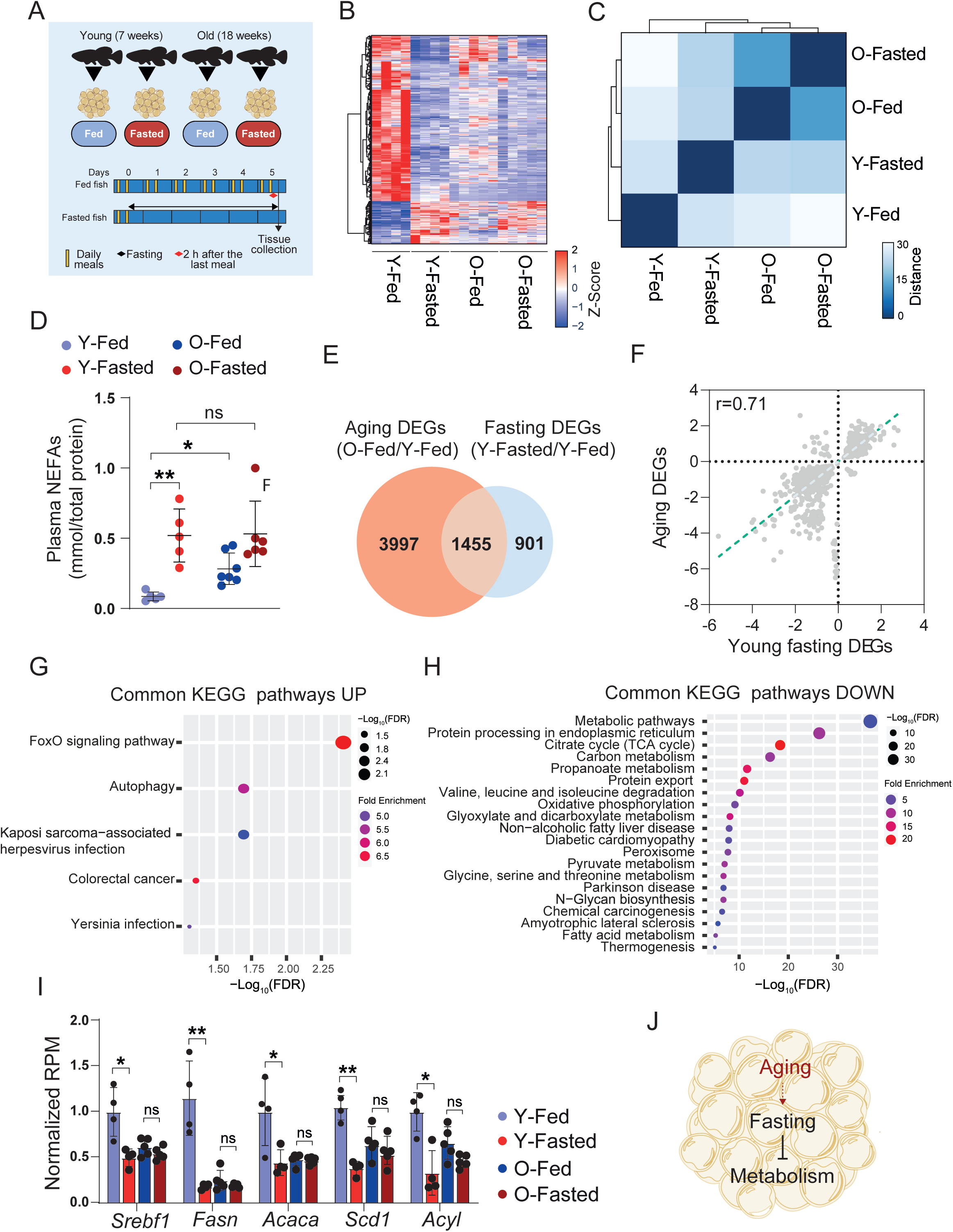
Aging alters the physiological response to fasting in the adipose tissue. **A**) Schematic of the food deprivation protocol. Control fish were fed twice daily (∼8:30 AM and ∼1:30 PM) and sacrificed 2 hours after the last meal. Food-deprived fish were fasted for five days and euthanized along with control fish. **B**) Hierarchical clustering of expression changes for fasting-induced genes (FDR<0.05). Colors represent the z-score range. **C** Hierarchical clustering of RNA-seq sample groups based on the Euclidean distance. **D**) Non-esterified fatty acids (NEFAs) quantification (mmol/plasma total protein quantification), 5-8 fish/group. **E**) Venn diagram showing the overlap between fasting and aging DEGs (Hypergeometric test: P=4.5e^-14^). **F**) Scatter plot of log2 fold changes for genes differentially expressed during aging and fasting (Pearson correlation: r=0.71, P<0.0001). The aging effect is depicted on the y-axis, and the fasting effect on the x-axis. **G-H**) KEGG pathway enrichment analysis of commonly up (G) and down-regulated genes (H). Q values are adjusted P-values using the false discovery rate procedure and are represented by a negative log10 scale (x-axis). **I**) Relative expression of DNL genes. Data values indicate RPM over the average RPM value of young fed samples. **J**) Schematic model showing the age-related changes in the killifish adipose tissue. Data in **D** and **I**) are presented as mean ± S.D. Significance was measured by Mann-Whitney U test in D and FDR in I. **\***:P<0.05; **\*\***: P<0.01; **\*\*\***:P<0.001.

To gain a more comprehensive understanding of the relationship between fasting and aging, we directly compared the transcriptional response of fasting (young fasted/young fed) with that of aging (old fed/young fed). Remarkably, this comparison showed an overlap of 1455 genes (22.9%, P=4.3e^-14^) (Figure 1E, supplementary tables 4 and 5). Furthermore, a large percentage (92%) of overlapping DEGs was altered in the same direction by fasting and aging (Figure 1F), indicating similar transcriptional regulation. KEGG enrichment analysis of the overlapping DEGs revealed a reduction of protein trafficking, energy production pathways (TCA-cycle and oxidative phosphorylation) (Figure 1G-1H, supplementary table 6), and several biosynthetic pathways, including the de novo lipogenesis pathway (DNL) (Figure 1I), crucial to maintaining adipogenesis. Thus, adipose tissue adopts a fasting-like transcriptional program (FLTP) that represses energy and biosynthetic metabolism in older fish (Figure 1J).

Looking at DEGs exclusive to old individuals after food deprivation, we observed that 4 of the top 10 up-regulated genes were implicated in the innate immune response (Figure 2A, supplementary table 3). Notably, immunity-related genes appeared significantly upregulated upon fasting only in older animals (Figure 2B), suggesting the presence of an age-related inflammatory signature associated with fasting. In addition, standard histology revealed a high number of infiltrating cells in the adipose tissue, skeletal muscle, and heart in old fasted animals (Figure 2C). As such, we stained for L-plastin, a pan-leukocyte marker in fish, revealing that old fasted fish showed a higher presence of immune-reactive cells in such organs (Figure 2C-2F). Notably, L-plastin+ reactive cells mostly surrounded adipocytes (crown-like structures) or, in some cases, skeletal muscle fibers. Thus, the physiological response to food deprivation in the adipose tissue is associated with an enhanced inflammatory signature in old animals.

**Figure 2:**
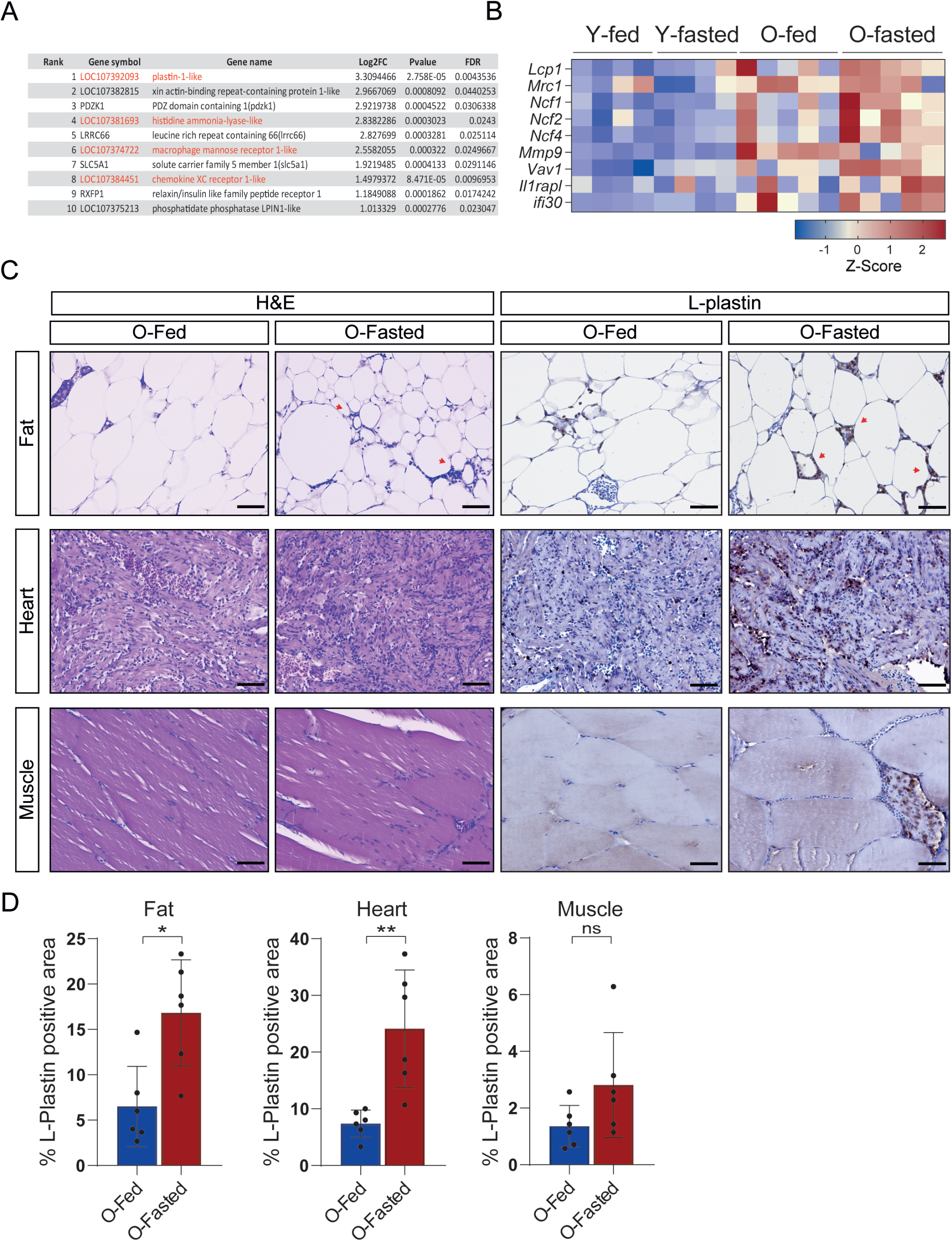
Fasting is associated with immune response in older animals. **A**) Top 10 fasting-induced genes in old fasted compared to old fed fish. Immune-related genes are indicated in red. **B**) Z-score heatmap of immune response-associated genes across all the samples. **C**) H&E staining (on the left) and L-plastin immunostaining (on the right). Red arrowheads indicate crown-like structures in the adipose tissue. **D**) Quantification of the area occupied by L-plastin positive cells over the total analyzed area, n=6 fish/group. Scale bar=100µm. Data in **D**) are presented as mean ± S.D. Significance was measured by Mann-Whitney U test. **\***:P<0.05; **\*\***: P<0.01;

FLTP and the observed reduction of biosynthetic pathways indicated a possible disruption of anabolic signals in older animals. The energy sensing pathway AMPK comprises seven different subunits (two α, two β, three γ) that assemble to form up to 12 heterotrimeric complexes. The physiologic relevance of all these different complexes is mainly unknown. Notably, fasting induced a significant downregulation of the AMPK γ1 subunit encoding gene *Prkag1* and upregulation of the γ2 encoding gene *Prkag2* at the transcriptional level (Figure 3A). This pattern was visible in old animals, whether fed or fasted, suggesting a possible dysregulation of AMPK signaling. Recent studies identified rare variants of specific AMPK γ subunits associated with human longevity^18^. Thus, we wished further to explore the potential connection between nutritional state and AMPK γ subunits expression. As we had initially observed regulation of the γ subunits upon five days of food deprivation, we next sought to determine their expression upon shorter fasting regimes and refeeding. Five groups of young killifish were fasted either for 0,18, 24, 48, 72 hours, and another two groups fasted for 72 hours and subsequently re-fed either for 6 and 24 hours, respectively. Remarkably, *Prkag1* expression was downregulated within 18 hours after food deprivation and appeared downregulated in all fasted groups, but rapidly increased within 6 hours of refeeding to level off within 24 hours of refeeding (Figure 3B). Conversely, *Prkag2* expression increased at 18 hours of fasting, peaked at 48 hours, and rapidly decreased with refeeding (Figure 3B). Extending the analysis to other organs, we observed a similar expression pattern in skeletal muscle, liver, and intestine (Extended Data Figure 2A-2C). Thus, our findings revealed that AMPK γ subunits exhibit an inverse oscillatory expression in response to the nutritional state. Next, we evaluated the effect of aging on γ-subunit expression dynamics. In this case, we monitored the expression of γ subunits at 0 and 48h of fasting, and 48h of fasting followed by 24h of refeeding. Surprisingly, the expression of γ-subunits in most of the analyzed tissues was not restored to a steady state upon refeeding in old animals (Figure 3C; Extended Data Figure 2D-2F). Thus, FLTP associates with a dysregulated expression of AMPK γ subunits in old animals.

**Figure 3:**
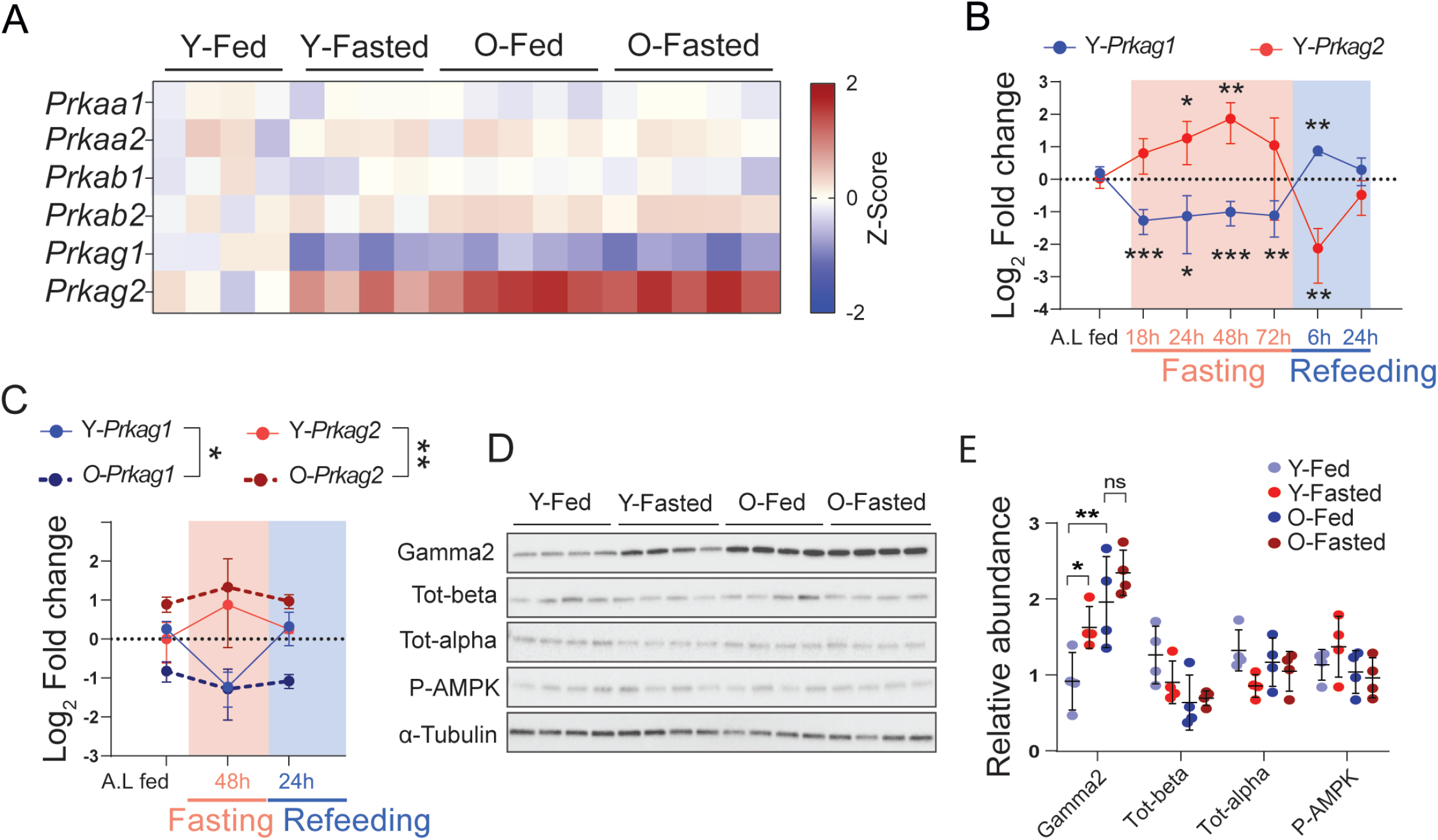
Fasting and aging modulate the expression of AMPK γ subunits. **A**) Z-score heatmap of AMPK subunits gene expression across all the samples. **B**) Log_2_ fold change of *Prkag1* and *Prkag2* relative expression upon fasting (0, 18, 24, 48, and 72 hours) and refeeding (72 hours of fasting followed by 6 and 24 hours of refeeding) in 7 weeks old fish, n=4/5 fish/group. Data values indicate fold change over the average value of “ad libitum” fed control animals (A.L). **C**) Log2 fold change of *Prkag1* and *Prkag2* relative expression upon fasting (48h) and refeeding (48h of fasting followed by 24 hours of refeeding) in young (7 weeks old, solid lines) and old individuals (18 weeks old, dashed lines), n=4/group. Data values indicate fold change over the average value of young fed control animals (A.L). **D**) Representative immunoblot of γ2 subunit, total-alpha, total-beta, phospho-Thr172-AMPKα, and α-tubulin. **E**) Quantification of protein abundance by densitometric analysis n=4/group; each dot represents an independent biological replicate. All data in **B, C, E**) are presented as mean ± S.D. Significance was obtained by one-way ANOVA followed by Tukey’s post-hoc test in **B** and **E)**, two-way ANOVA followed by Sidak multiple comparison test in **C**). **\***:P<0.05; **\*\***: P<0.01; **\*\*\***:P<0.001.

### Sustained activity of the AMPK_γ1_ complex prevents age-associated FLTP

As the canonical function of γ subunits is to activate AMPK by allosteric regulation, we tested whether changes in the γ subunit’s stoichiometry influence AMPK activity in response to energy depletion. Using cross-reactive antibodies to γ2, we confirmed protein expression to be induced upon fasting and constitutively expressed in old fed or fasted fish while detecting no changes in other AMPK subunits nor phospho-AMPK levels (Figure 3D-3E). However, we could not monitor γ1 protein levels because available antibodies did not cross-react (data not shown). Shifts in the γ-subunit expression did not influence phospho-AMPKα levels. This led us to speculate that the dynamic interchange of γ subunits might alter the function of AMPK according to the nutritional status. AMPK is a positive regulator of catabolism; thus, we considered it surprising that γ1 expression was induced by anabolic stimuli (refeeding) and repressed by catabolic stimuli (fasting). As older animals showed FLTP and reduced expression of γ1, we asked whether selective stimulation of the AMPK_γ1_ complex activity could counteract FLTP and restore the normal response to anabolic stimuli in older animals. The γ1 R70Q mutation suppresses the inhibitory properties of ATP and results in constitutive activation of AMPK_γ1_ complexes^19^. Given the high conservation of AMPK subunits across killifish and mammals (Extended data Figure 3A), we introduced the (R70Q) mutation into the endogenous γ1 locus by CRISPR genome engineering (referred to as γ_1(R70Q),_ Figure 4A) and, as γ1 expression is suppressed during aging, we generated another line overexpressing constitutively activated γ1 subunits bearing the (R70Q) mutation (referred to as Ubi:γ_1(R70Q)_; Figure 4B; see Materials and Methods and extended data Figure 7 for details). We then determined the phosphorylation levels of the AMPKα subunit residue at Thr172, required for AMPK activation, and its downstream substrate ACC in the adipose tissue of middle-age wild-type, Ubi:γ_1(R70Q),_ and γ_1(R70Q)_ fish. Ubi:γ_1(R70Q)_ line only showed elevated phospho-AMPKα levels, whereas both mutant lines showed elevated P-ACC levels, although the effect appeared stronger in Ubi:γ_1(R70Q)_ (Extended data Figure 3B-4C). Thus, in line with mammalian data, the introduction of the R70Q mutation resulted in chronic activation of the AMPK_γ1_ complex. We next determined the physiologic response to food deprivation in fish with increased AMPK_γ1_ activity, as we did previously with wild-type animals (Figure 4C). For this experiment, we employed the Ubi:γ_1(R70Q)_ line as it showed a more robust AMPK_γ1_ complex activation. RNA-seq analysis identified 3185 DEGs comparing young fasted/young fed and, surprisingly, 5465 DEGs comparing old fasted/old fed (Extended data Figure 3D-4E, supplementary table 7 and 8). Despite the high number of DEGs induced by fasting, KEGG pathway analysis revealed that the differentially regulated processes in Ubi:γ_1(R70Q)_ were consistent with those observed in young wild-type (Extended data Figure 3F). These data indicate that sustained AMPK_γ1_ retains a more youthful fasting response in older fish. Next, we assessed whether this was due to the absence of FLTP in old Ubi:γ_1(R70Q)_ fish. Unlike wild-type animals, the normalized expression of fasting-responsive genes (FDR<0,05) in old mutant fed fish more closely resembled the profile of fed fish (Figure 4D), and coherently, distance matrix analysis revealed young and old fully fed groups clustering together (Extended data Figure 3G). Further, aging and fasting transcriptional responses overlapped for only 398 genes (9%) (Figure 4E, supplementary table 9). Of those, 36% showed an opposite regulation (Figure 4F). Evaluating the age-related transcriptional changes in Ubi:γ_1(R70Q)_ fish (old fed/ young fed), we observed little or no sign of reduced energy and biosynthetic pathways (Extended data Figure 3H). In line with this, neither Ubi:γ_1(R70Q)_ nor γ_1(R70Q)_ fish showed age-related reduced expression of DNL genes (Figure 4G, Extended data Figure 3I), indicating these mutant lines share a similar transcriptional profile. Altogether these data indicate that sustained AMPK_γ1_ complex activation counters the age-associated FLTP and maintains energy and anabolic metabolism late in life.

**Figure 4:**
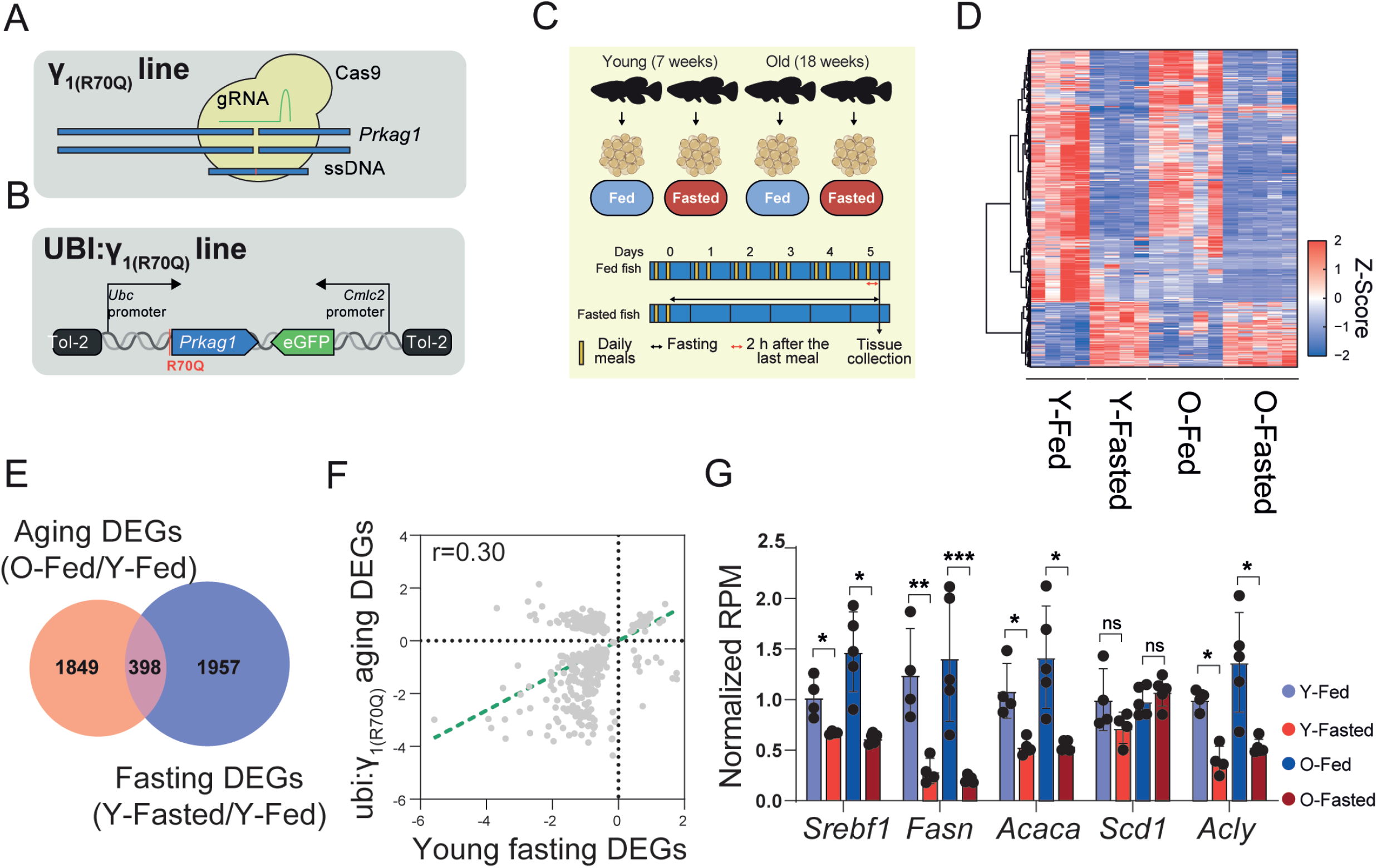
Sustained activation of AMPK_γ1_ prevents FLTP in old fish. **A-B**) Schematic illustration showing the CRISPR/Cas9 line generation (Y_1(R70Q)_) in A, and the Tol-2 line (Ubi:_Y1(R70Q)_) in B. **C**) Schematic of the food deprivation protocol. Control fish were fed twice daily (∼8:30 AM and ∼1:30 PM) and sacrificed 2 hours after the last meal. Fasted fish were fasted for five days and euthanized along with control fish. **D**) Hierarchical clustering of expression changes for fasting-induced genes (FDR<0.05). Colors represent the z-score range. **E**) Venn diagram showing the overlap between fasting and aging DEGs (Hypergeometric test: P=0.99). **F**) Scatter plot of log_2_ fold changes for genes differentially expressed during aging and fasting (Pearson correlation: r=0.30, P<0.001). The aging effect is depicted on the y-axis, and the fasting effect on the x-axis. **G**) Relative expression of DNL genes. Data values indicate RPM over the average RPM value of young fed samples. Data **G**) are presented as mean ± S.D, and significance was measured by FDR in I. **\***:P<0.05; **\*\***: P<0.01; **\*\*\***:P<0.001.

### Chronic activation of the AMPK_γ1_ complex sustains anabolism and regenerative processes late in life

Refeeding stimulates higher energy expenditure, protein synthesis, and cellular proliferation, all processes required for tissue turnover and repair but commonly repressed during aging. Thus, we asked whether the sustained anabolic processes induced by AMPK_γ1_ implied higher regenerative processes in older mutant fish. To this end, we compared the transcriptome profile of old fed Ubi:γ_1(R70Q)_ to that of old wild-type fish. We identified 2963 differentially expressed genes (FDR<0.05, supplementary table 10). KEGG pathway analysis revealed that the strongest upregulated categories were proteostasis, DNA replication, and repair (Figure 5A, supplementary table 11). Within these categories, we found upregulated genes involved in DNA synthesis, DNA double-strand break repair (DSBR), including *Rad51, Xrcc1*-*2*, and *Atm*, as well as many genes encoding proteasomal subunits (Figure 5C), suggesting increased cellular proliferation, improved DNA damage surveillance and proteome turnover. Furthermore, upregulated genes were enriched for processes such as carbon metabolism and TCA cycle (Figure 5A, supplementary table 11), indicative of increased energy expenditure. In line with this, we found upregulation of multiple genes coding for cytosolic and mitochondrial ribosomes and several OXPHOS complex subunits. On the other hand, down-regulated genes were enriched for insulin secretion, adipocytokine signaling, and inflammatory processes (Figure 5B, supplementary table 12). Here were found many genes whose expression positively correlates with obesity, insulin resistance (IR), type-2 diabetes (T2D), and chronic inflammation, such as *Irs2, Angptl4, Lepr, Camkk2*, and *Tbk1* (Extended data Figure 4A). Adipogenesis, including the new production of adipocytes, dramatically declines with aging promoting lipodystrophy and metabolic dysfunction^20^. As transcriptomic analysis indicated a possible signature of increased cellular proliferation in Ubi:γ_1(R70Q)_, we performed a pulse-chase EDU labeling experiment to determine the proliferative status of the adipose tissue in older animals. Notably, both Ubi:γ_1(R70Q)_ and γ1_(R70Q)_ lines showed a higher number of dividing cells compared to age-matched wild-type fish (Figure 5D-5E). Overall, these data indicate that increased AMPK_γ1_ complex activity sustained proliferative processes in the adipose tissue of older animals.

**Figure 5:**
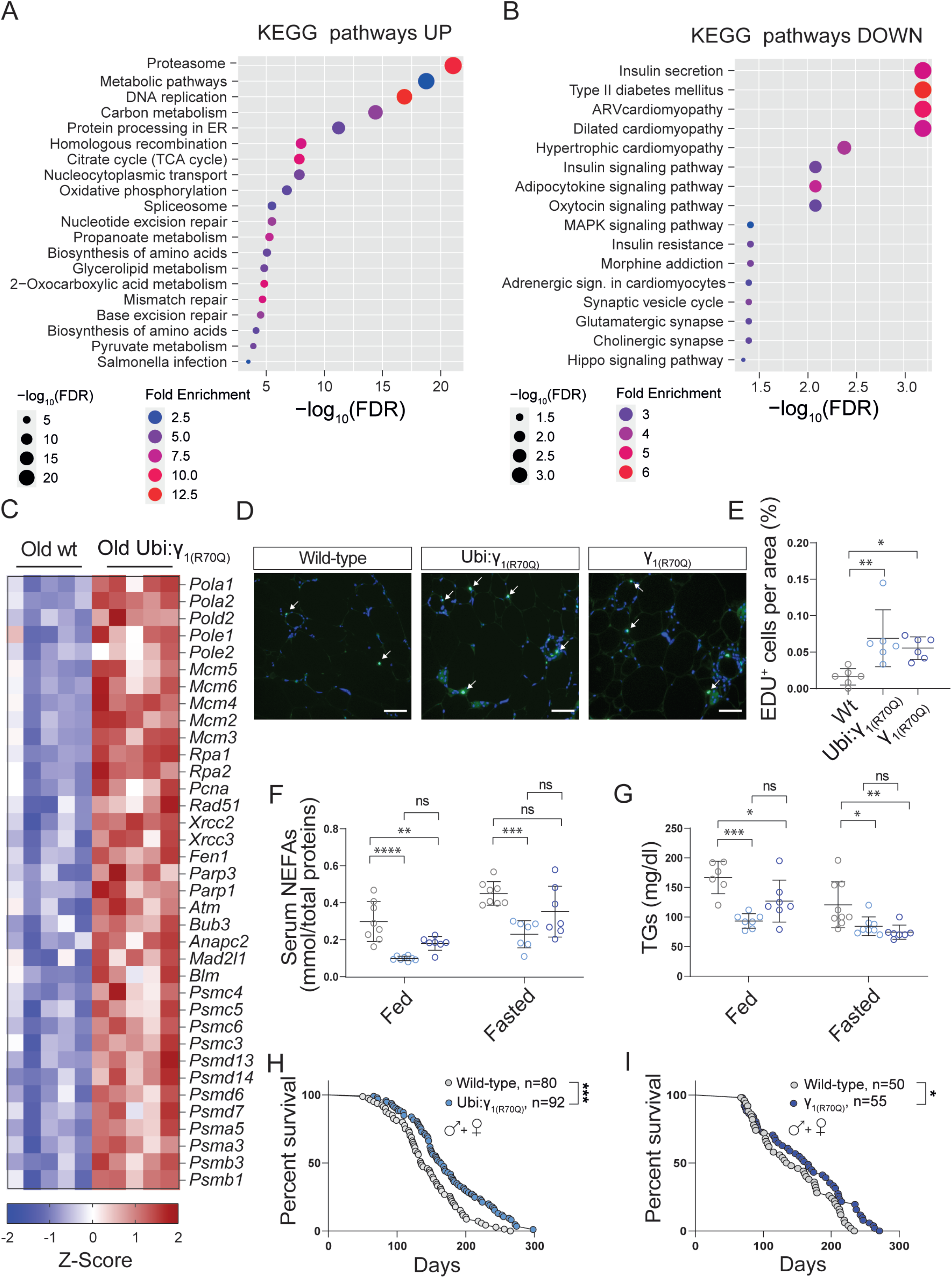
Sustained activation of the AMPK_γ1_ complex promotes metabolic health and longevity. **A-B**) KEGG pathway enrichment analysis of up (A) and down-regulated genes (B) in old Ubi:γ_1(R70Q)_ compared to old wild-type fish under the normal fed condition. Q values are the adjusted P-values using the false discovery rate procedure and are represented by a negative log10 scale (x-axis). **C**) Z-score heatmap showing differentially expressed genes involved in DNA synthesis and repair, and proteostasis. **D**-**E**) Representative images and relative quantification of EDU-labeled cells in the adipose tissue of 18 weeks old wild-type and mutant lines. Scale bar=100µm. White arrows indicate EDU-positive cells. **F**) Non-esterified fatty acids (NEFAs) quantification (mmol/plasma total protein quantification) in fed and fasted (48h) 18 weeks old fish, 6-8 fish/group. **G**) Blood triglycerides quantification (mg/dl) in fed and 48 hours-fasted 18 weeks old fish, 6-8 fish/group. **H**-**I**) Kaplan-Meier survival analysis of Ubi:γ_1(R70Q)_ and Y_1(R70Q)_ lines, sex combined. All data in **E-G**) are presented as mean ± S.D. Significance was obtained by one-way ANOVA followed by Tukey’s post-hoc test in **E-G**), and by two-tailed log-rank calculation in **H-I**). **\***:P<0.05; **\*\***: P<0.01; **\*\*\***:P<0.001.

### Chronic activation of the AMPK_γ1_ complex promotes metabolic health and longevity

Next, we examined the health parameters of these lines. Old Ubi:γ_1(R70Q)_ and γ_1(R70Q)_ fish showed a slight reduction in body weight compared to age-matched wild-type fish, though the latter did not reach statistical significance (Extended Data Figure 4B). Notably, the food consumption of 18 weeks old mutant fish monitored over one week was not significantly different compared to old wild-type fish (Extended Data Figure 4C). Body scan analysis (% of volume, extended Data Figure 4D) and visceral fat quantification (% of body weight; extended Data Figure 4E) revealed a general reduction of visceral adiposity in both sexes, while subcutaneous adiposity decreased only in males. As females allocate a substantial amount of lipids into eggs, we wondered whether the reduction in adiposity could impact egg lipid content. However, we saw comparable levels of egg lipid content between mutant and wild-type fish (Extended Data Figure 4F). In addition, in line with higher expression of DNL genes and proliferative potential, Ubi:γ_1(R70Q)_ and γ_1(R70Q)_ displayed a higher number of small adipocytes and a reduced number of hypertrophic adipocytes relative to wild-type fish (Extended Data Figure 4G-4H). Interestingly, both Ubi:γ_1(R70Q)_ and γ_1(R70Q)_ lines showed reduced serum triglycerides (TGs) and circulating NEFAs levels under basal and fasting conditions, an effect that was stronger in Ubi:γ_1(R70Q)_ (Figure 5F-5G). They also showed reduced fasting blood glucose levels (Extended Data Figure 4I). Our findings suggest that chronic activation of the AMPK_γ1_ complex significantly improves lipid and glucose parameters in old killifish. Finally, we determined the effect of sustained AMPK_γ1_ activation on life span. As no gender gap in life expectancy was observed in single-housed killifish (Extended Data Figure 5A), we used both sexes for survival analysis. The median survival of Ubi:γ_1(R70Q)_ and γ_1(R70Q)_ (sexes combined) was 20% and 14% relative to wild-type fish, respectively (Figure 5H-5I). Analysis of each sex separately showed a 19.1% and 8% (not statistically significant) increase in males and 21.6% and 15.8% increase in females media lifespan of Ubi:γ_1(R70Q)_ and γ_1(R70Q)_ respectively (Extended data Figure 5B-5E). Thus, our data reveal that AMPK_γ1_ activity is induced by refeeding to stimulate regenerative processes, thus promoting health and longevity.

### *Prkag1* expression is a marker of healthy aging in humans

Finally, we wished to evaluate the relevance of AMPK_γ1_ for human health and longevity. Defour et al. performed a transcriptome analysis of human subcutaneous adipose tissue (SAT) in response to fasting. They reported modulation of the AMPK γ2 subunit^21^, potentially indicating the presence of a nutritional-dependent expression of γ subunits in humans. In light of these findings, we next used the human genotype tissue expression (GTEx) dataset to evaluate the expression of γ subunits as a function of age. In line with killifish data, *Prkag1* expression significantly decreased as a function of age in SAT, blood cells, heart in both sexes, and liver in males only (Figure 6A-6E). On the contrary, *Prkag2* expression significantly increased in SAT only (Extended data Figure 6A-6E). Blood showed one of the most substantial age-related *Prkag1* downregulation among analyzed tissues. Thus, we asked whether the presence of such downregulation in the blood of older humans could be associated with multimorbidity, frailty, and mortality risk. To this end, we determined the expression of *Prkag1* and *Prkag2* by qPCR in peripheral blood mononuclear cells (PBMCs) from 93 older donors (aged between 65 and 90 years). Study participants were patients hospitalized for non-neurological, non-surgical medical conditions and included solely according to age. Importantly, we established a cohort covering a broad range of age-related diseases with very heterogenous functional levels. For all participants, the multidimensional prognostic index (MPI) was recorded at the time of presentation to the hospital. The MPI is a validated prognostic model comprised of eight components, namely, Activities of Daily Living (ADL), Instrumental ADL (IADL), Short Portable Mental Status Questionnaire (SPMSQ), Cumulative Illness Rating Scale - Comorbidity Index (CIRS-CI), Mini Nutritional Assessment Short Form (MNA-SF), Exton Smith Scale (ESS), Number of medications (NM), and the quality of the social support network (Figure 6G). Thus, it is based on information regarding functional, psychosocial, clinical, and nutritional status^22^. The MPI has been shown to predict older individuals’ mortality rate more accurately than the classical physical frailty score^23^. Higher values of MPI reflect poor health outcomes and higher mortality risk^24^. We found that γ1, but not γ2, expression was inversely correlated with MPI score (Figure 6F, Extended data Figure 6F). However, we observed no correlation between γ subunits expression and the age of the donors (Extended data Figure 6G-6H), indicating that the relationship between MPI and γ1 expression is independent of the chronological age of the donors at this later stage of life. Looking at the individual MPI components, we observed a significant negative correlation of *Prkag1* expression with CIRS-CI and NM (Figure 6H-6I). At the same time, MNA, ADL, IADL, and ESS were positively correlated with *Prkag1* expression (Figure 6J-6O), indicating reduced multimorbidity and improved nutritional status and functional skills in older individuals to be associated with higher γ1 expression. On the contrary, *Prkag2* expression was not associated with any MPI component (Extended data Figure 6I-6P). This highlights the functional relevance of considering the individual γ subunits and indicates that changes in *Prkag1* expression are specific rather than a consequence of general changes in protein levels due to disease. These data suggest that *Prkag1* expression is a possible indicator of healthy human aging.

**Figure 6:**
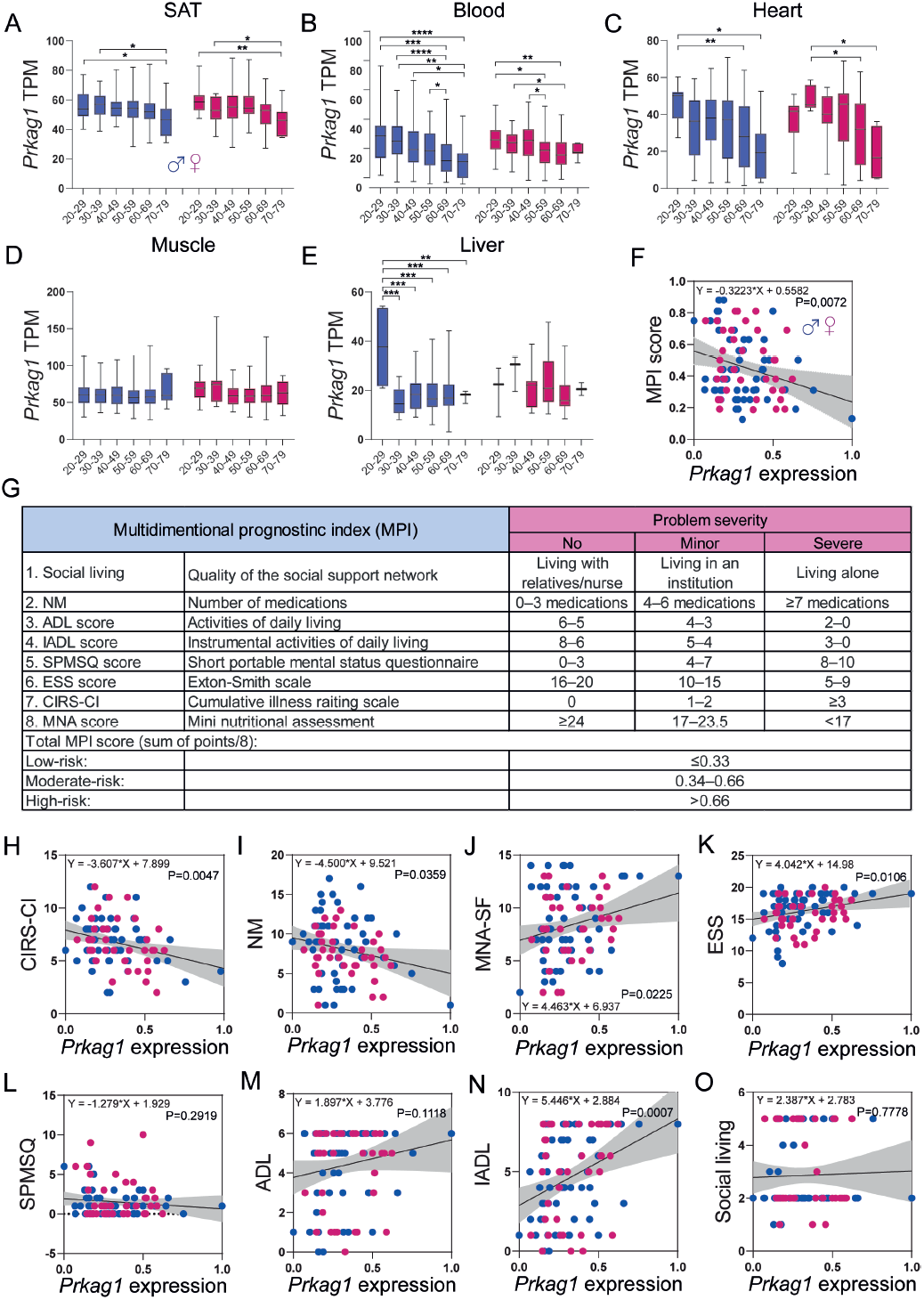
Human AMPK γ1 expression is a biomarker of healthy aging. **A**-**E**) Box plots showing human normalized *Prkag1* expression levels across age groups in decadal brackets. Data are available in the GTEx Consortium (GTEx analysis V8). **F**) Scatter plot showing the relationship between *Prkag1* expression and MPI score in males (blue dots) and females (pink dots). The black line represents a linear model fit with a 95% confidence interval in gray. **G**) Calculation of the multidimensional prognostic index (MPI). **H-O**) Scatter plots comparing the relationship between *Prkag1* expression and MPI-subdomains. Black lines represent linear model fit with a 95% confidence interval in gray. Significance was obtained by one-way ANOVA followed by Tukey’s post-hoc test in A-E. **\***:P<0,05; **\*\***: P<0,01; **\*\*\***:P<0,001; ****:P<0,0001.

## Discussion

In this study we determined the transcriptional response to food deprivation in visceral adipose tissue (VAT) of young and old fish. We found that aging promotes a fasting-like transcriptional program (FLTP) that reduces anabolic activities, thereby setting VAT in a quasi-quiescent state. However, because we only measured FLTP in VAT, we do not yet know the full extent of FLTP in other tissues. Notably, we observed dysregulation of AMPK γ subunits in adipose and other aging tissues, implying the presence of FLTP as a broad response. In line with our data, Montesano et al. reported the expression of the orexigenic neuropeptide Y (NPY), normally induced by fasting, to progressively increase in killifish diencephalon (thalamic area) during aging^25^. These data indicate that FLTP might not be limited to the adipose and could reflect intrinsic tissue processes or a systemic effect of FLTP in adipose on other tissues.

Another critical question is whether FLTP occurs in higher vertebrate species. Human aging is often accompanied by a significant reduction of subcutaneous adiposity, which progressively relocates into visceral depots^20^. Loss of SAT associates with poor health outcomes in humans, yet the reasons remain unknown. In line with FLTP observed in killifish, age-related changes in human SAT include suppression of energy metabolism, adipogenesis^20,26^, and an increased number of infiltrating immune cells^27^. In addition, older individuals often exhibit elevated plasma NEFAs^28^ and increased liver steatosis^29^, both signs of an ongoing fasting response. Finally, we could show a broad age-associated reduction of γ1 expression in human SAT. Altogether these findings suggest the possible presence of FLTP in human SAT, which could reflect the suppression of self-renewal processes and promotion of elevated plasma lipids species. Another tissue where FLTP might take place is skeletal muscle since age-related sarcopenia is characterized by the inability of the muscle to maintain the correct protein synthesis in response to feeding or exercise, a condition defined as anabolic resistance^30^. Thus, multiple lines of evidence support the notion that aging is associated with a reduction of anabolism across vertebrate species.

Aging is characterized by an overall decline of the regenerative capacities of the body^31,32^. Remarkably, we observed that old Ubi:_γ1(R70Q)_ or γ1(R70Q) fish exhibit a sustained response to anabolic stimuli in old age, higher cellular proliferation, metabolic health, and longer life compared to wild-type animals. While AMPK is best known as a regulator of catabolic processes, these data suggest that maintaining the proper balance of anabolism and catabolism late in life could have a rejuvenating effect and stimulate regenerative processes. Accordingly, transfection of Yamanaka factors, which by definition are promoters of growth and cellular proliferation, counters aging and promotes rejuvenation in older animals^33-36^. These findings apparently contrast with the anti-aging effects of mTOR inhibition in middle-aged mice^37,38^. However, such effects most likely arise from an enhanced activation of autophagy rather than a reduction of anabolism^39^. Notably, the global protein translation rate is not altered in S6K-deficient muscle cells^40^, while the overexpression of constitutively active S6K does not abolish lifespan extension by chronic or short-term rapamycin treatment in flies^39^. It is also possible to speculate that the reduction of mTOR could trigger feedback activation of other pro-anabolic regulators. In this regard, S6K-/-mice and worms were shown to have increased AMPK activity, and such activation was required for S6K-/-mediated longevity^41^. Thus, strategies aimed at sustaining anabolic metabolism balance and tempering catabolism late in life appear successful in promoting tissue rejuvenation.

The evidence that anabolic signals become impaired in older individuals could have important implications for anti-aging interventions such as dietary restriction (DR) or intermittent fasting (IF). In fact, the protective effects mediated by either DR or IF derive from the coupled effects of fasting and refeeding that constrain metabolism to shift from catabolism to anabolism periodically. The evidence that old animals become refractory to the refeeding arm implies that DIs initiated late in life might exacerbate catabolic activities, fail to reinstate the proper anabolic profile upon refeeding, and ultimately promote tissue wasting. In line with this idea, we showed that old individuals, as opposed to young, displayed an enhanced inflammatory signature upon food deprivation.

The evidence that the AMPK complex dynamically changes γ subunits according to the nutritional status highlights the importance of complex composition as a new layer of regulation in defining AMPK functions and should be considered for further studies on AMPK physiology. Classically, AMPK is inhibited by high ATP levels and activated by increased AMP/ATP ratio or low-glucose concentration^42,43^. However, the evidence that γ1 expression is low under fasting and high under food availability indicates that this subunit allows AMPK to function under high-energy conditions. Notably, among all the γ subunits, γ1 only is sensitive to ADP, whose levels are considerably higher compared to AMP under normal feeding conditions^44^. By contrast, the γ2 subunit appears insensitive to ADP stimulation but is the most responsive to AMP fluctuations^43^. Consistently we show that the AMPK γ2 subunit is induced under fasting conditions in young healthy fish. This could indicate that AMPK_γ2_ function is physiologically required exclusively upon fasting and explain why, unlike AMPK_γ1_, chronic activation of AMPK_γ2_ leads to metabolic dysfunctions in normally fed mice or humans^45,46^. Astre et al. showed that mutation in the APRT gene, a key enzyme in AMP biosynthesis, mimics a caloric-restriction effect, inducing lifespan extension in killifish males^47^. This effect was associated with higher expression of the γ2 subunit, possibly indicating that higher activation of AMPK_γ2_ is beneficial under energetic stress conditions and implies that various dietary interventions might represent a concrete strategy to counter the pathological effect of AMPK_γ2_ hyperactivity in humans.

Finally, we found that PBMCs γ1 expression in humans anticorrelates, independent of chronological age, with multidimensional frailty as measured by the MPI score as well as with its nutritional, functional, and multimorbidity subdomains. This observation strongly suggests this gene as a possible hallmark of healthy aging and human robustness. A large body of recent evidence shows that multidimensional frailty, but not chronological age, is associated with adverse outcomes with increasing age. While the MPI measures multidimensional frailty with the highest clinimetric properties, no biomolecular/genetic signature for it has been identified so far. Importantly, the evidence that γ1 but not γ2 expression inversely correlates with the MPI highlights the relevance of the specific AMPK complex composition and could have important therapeutic implications. In fact, AMPK-activating compounds are identified based on their ability to stimulate the kinase activity, but very often, without considering the nature of the complex activated. Thus, further studies are required to identify AMPKγ1-selective activating compounds and test their effect on the aging, robustness, and longevity of vertebrate species.

## Acknowledgments

We thank all members of the Antebi lab for scientific input, the bioinformatic core facility (Jorge Boucas, Franziska Metge) for help with data analysis, the phenotyping core facility (Andrea Mesaros and Martin Purrio) for microCT-scan analysis, the MPI-AGE FACS & Imaging Core Facility (Christian Kukat, Alexandra Just, and Marcel Kirchner) for support with histological preparations and microscopy. Raymond Laboy Morales for the help with Figures editing, Mattias Werres and Christina Paetzold for help with killifish husbandry, and Orsolya Symmons for critical input. We thank Jasmin Garha and Cornelia Böhme for excellent support with human biosampling. This work was funded by the Max Planck Society and the Deutsche Forschungsgemeinschaft (DFG, German Research Foundation) under Germany’s Excellence Strategy – CECAD, EXC 2030 – 390661388.

## Author contributions

R.R and A.A conceived the study, planned the experiments, and wrote the manuscript. R.R performed all the experiments with input from E.B, and C.C. A.B. and C.L. A.M., R-U.M., and M.C.P. provided human blood samples and relative MPI score. J.S. isolated hPBMCs. N.H. gave technical assistance. L.D. assisted with the fish hatching and breeding. All authors commented on the manuscript.

## Figure legends

**Extended Data Figure 1.**
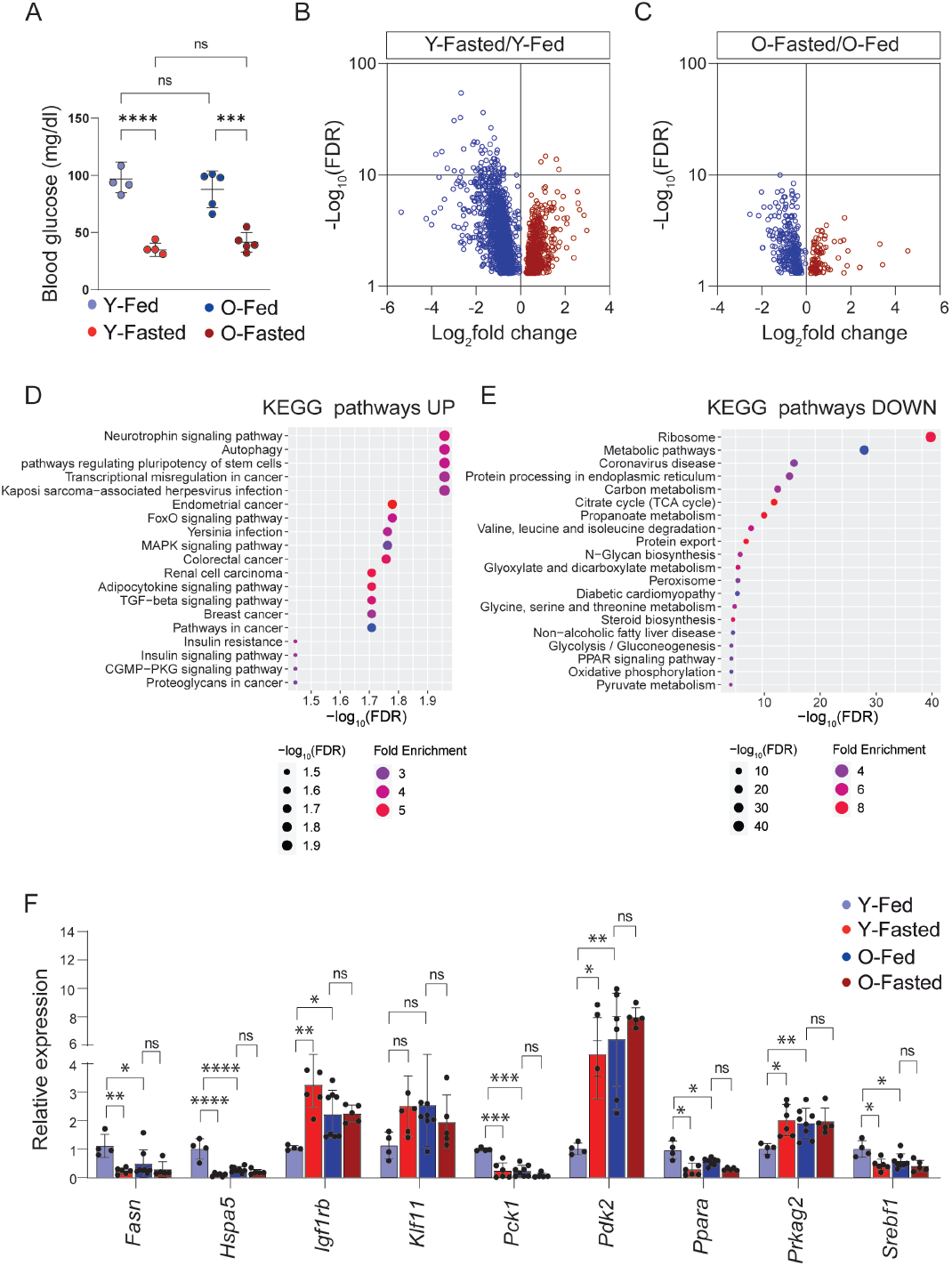
Aging associates with a fasting-like transcriptional program (FLTP). **A**) Blood glucose measurements of young and old fish under feeding or fasting regime, n=4-5 fish/group. **B**-**C**) Volcano plots showing differentially expressed genes (P<0,05) in young (B) and old (C) wild-type fish under fasting. **D**-**E**) KEGG pathway enrichment analysis of up (A) and down-regulated genes (B) in young fasted/fed fish. Q values are the adjusted P-values using the false discovery rate procedure and are represented by a negative log10 scale (x-axis). **F**) qPCR analysis of top fasting-regulated genes in young and old fish, 4-8 fish/group. Data values indicate fold change over the average value of young samples. Data in **A** and **F**) are presented as mean ± S.D. Significance was obtained by two-way ANOVA followed by Sidak multiple comparison test in A, by one-way ANOVA followed by Tukey’s post-hoc test in F. **\***:P<0,05; **\*\***: P<0,01; **\*\*\***:P<0,001.

**Extended Data Figure 2.**
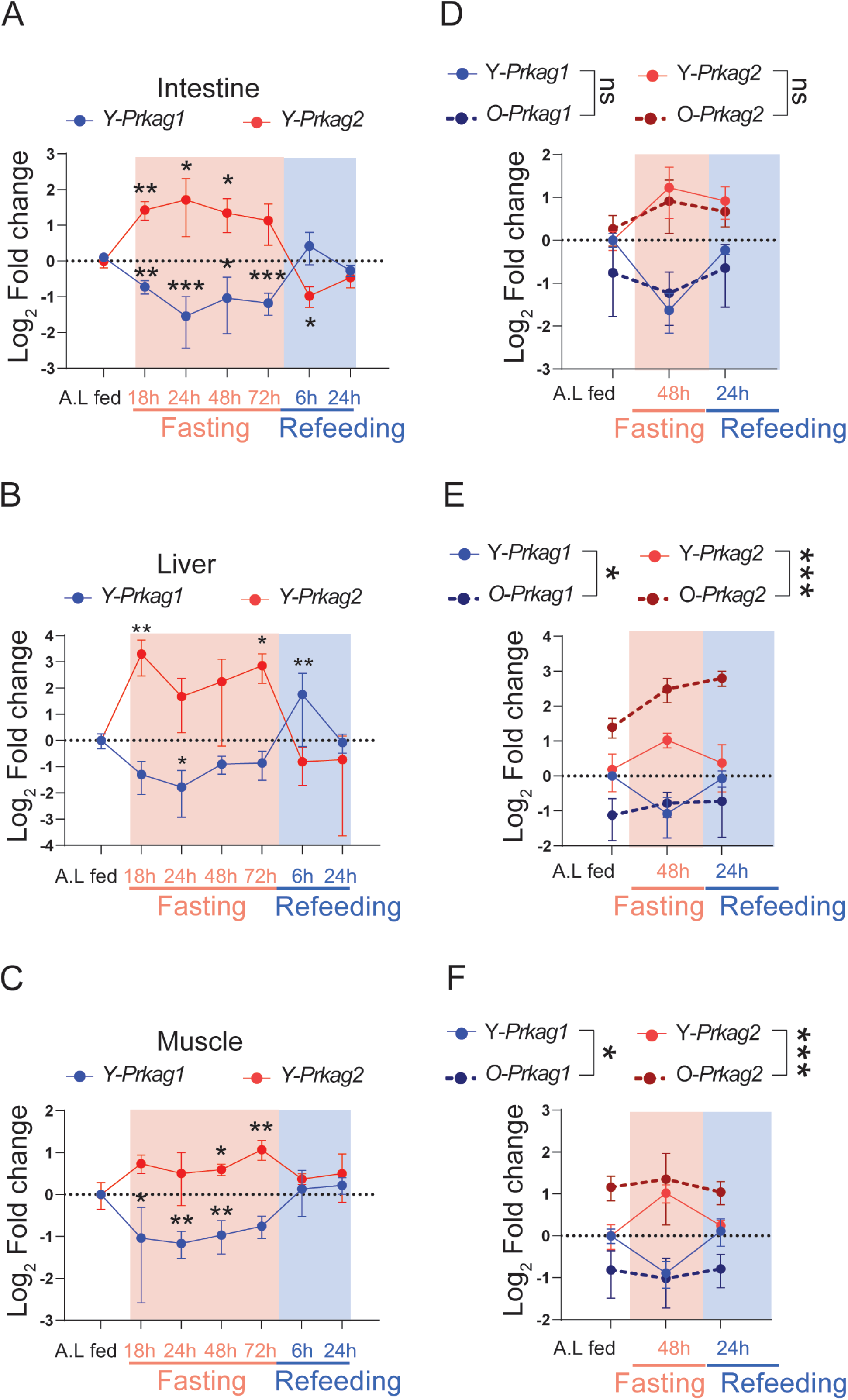
Nutritional-dependent regulation of AMPK γ subunits mRNAs in different tissues. **A-C**) Log2 fold change of *Prkag1* and *Prkag2* relative expression upon fasting (0, 18, 24, 48, and 72 hours) and refeeding (72 hours of fasting followed by 6 and 24 hours of refeeding) in the intestine, muscle, and liver in 7 weeks old fish, n=4/5 fish/group. Data values indicate fold change over the average value of “ad libitum” fed control animals (A.L). **D-F**) Log2 fold change of *Prkag1* and *Prkag2* relative expression upon fasting (48h) and refeeding (48h of fasting followed by 24 hours of refeeding) in young (7 weeks old, solid lines) and old individuals (18 weeks old, dashed lines), n=4 fish/group. Data values indicate fold change over the average value of young fed control animals (A.L). All the data in **A-F**) are presented as mean ± S.D data. Significance was obtained by one-way ANOVA followed by Tukey’s post-hoc test in **A**-**C**), two-way ANOVA followed by Sidak multiple comparison test in **D**-**F**) **\***:P<0,05; **\*\***: P<0,01; **\*\*\***:P<0,001.

**Extended Data Figure 3.**
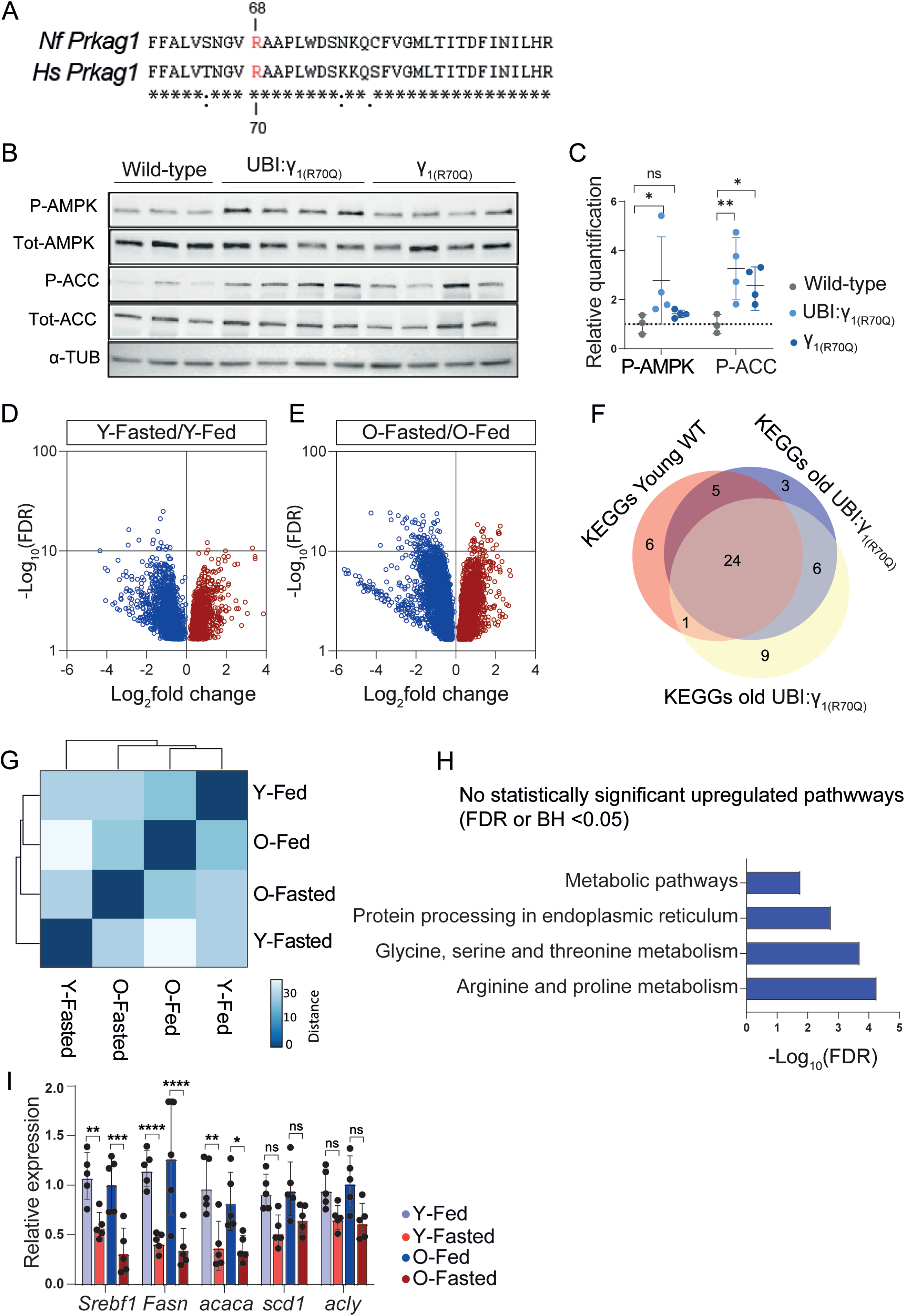
Sustained activation of the AMPK_γ1_ complex prevents FLTP. **A**) Evolutionary conserved AMPK γ1 amino acid sequence between humans and killifish. Red highlights the critical amino acid residue R70 (R68 in killifish). **B-C**) Representative Western blots and densitometric analysis of phospho-Thr172-AMPKα, total-AMPKα, phospho-S79-ACC, total-ACC, in 18 weeks old killifish adipose tissue. α-tubulin was used as a loading control. **D**-**E**) Volcano plots showing differentially expressed genes (FDR<0,05) in young (D) and old (E) Ubi:γ_1(R70Q)_ fish. **F**) Venn diagram depicting the overlap of KEGG pathways differentially regulated by fasting in young wild-type, young Ubi:γ_1(R70Q),_ old Ubi:Y_1(R70Q)_ fish (hypergeometric test P=1,2^e-12^). **G**) Hierarchical clustering of RNA-seq samples based on the Euclidean distance. **H**) KEGG pathway enrichment analysis of differentially regulated genes comparing old to young fed Ubi:γ_1(R70Q)_ fish. Q values are adjusted P-values using the false discovery rate procedure and are represented by a negative log10 scale (x-axis). **I**) qPCR analysis of DNL genes in young and old fed and fasted Y_1(R70Q)_ fish, n=5 fish/group. Data in **C** and **I**) are presented as mean ± S.D. Significance was obtained by one-way ANOVA followed by Tukey’s post-hoc test in C and I. **\***:P<0.05; **\*\***: P<0.01; **\*\*\***:P<0.001.

**Extended Data Figure 4.**
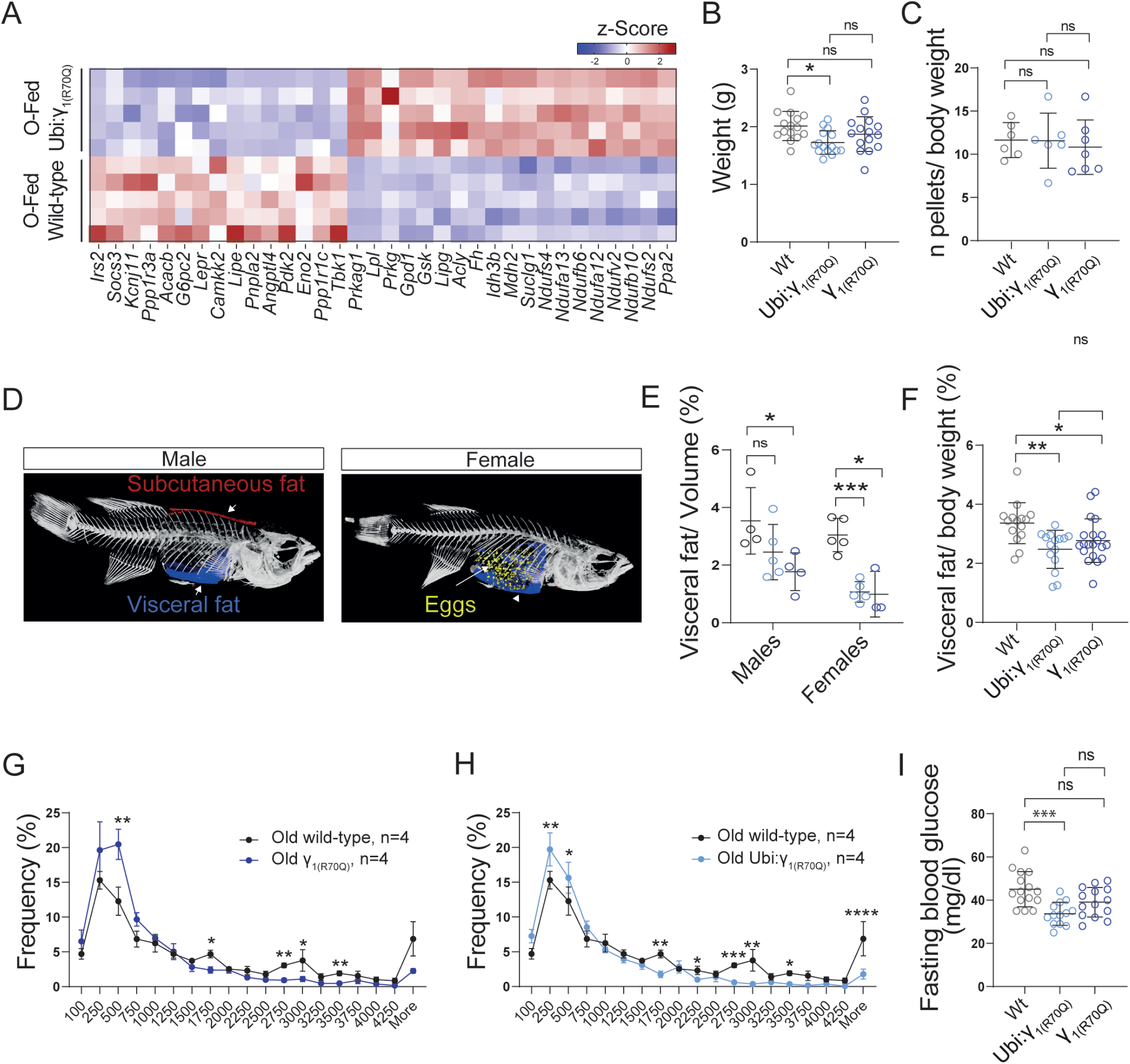
Sustained activation of AMPK_γ1_ promotes metabolic health and longevity. **A**) z-score heatmap of genes involved in lipid and glucose handling. **B**) Body weight comparison in18 week old wild-type, Ubi:γ_1(R70Q)_, and Y_1(R70Q)_ fish. **C**) Food consumption monitored over a week in 18 weeks old fish. Each dot represents the average value of pellets eaten/ body weight by each fish, n=6-7 fish/group. **D**) 3D Micro-CT analysis of whole-body composition in wild-type killifish. Visceral fat is indicated in blue, subcutaneous fat in red, and eggs in yellow. **E**) Micro-CT scan quantification of visceral fat in 18 weeks old fed wild-type and mutant lines (both sexes), n= 3-5 fish/group. **F**) Visceral fat quantification (milligrams of fat /body weight) of 18 weeks old wild-type (n=12), Ubi:γ_1(R70Q)_ (n=15) and Y_1(R70Q)_ (n=20) male fish. **G**-**H**) Frequency distribution of adipocyte area in 18-week-old wild-type, Ubi:γ_1(R70Q)_, and Y_1(R70Q)_ fish, n=4 fish/group. **I**) Fasting (48 hours) blood glucose levels (mg/dl) of 18 weeks old negative transgenics (n=15), Ubi:γ_1(R70Q)_ (n=12) and Y_1(R70Q)_ (n=14) male fish. Data in **B**-**I**) are presented as mean ± S.D. Significance was obtained by one-way ANOVA followed by Tukey’s post-hoc test in B-C-F-I, and by t-test in G-H. **\***:P<0,05; **\*\***: P<0,01; **\*\*\***:P<0,001.

**Extended Data Figure 5.**
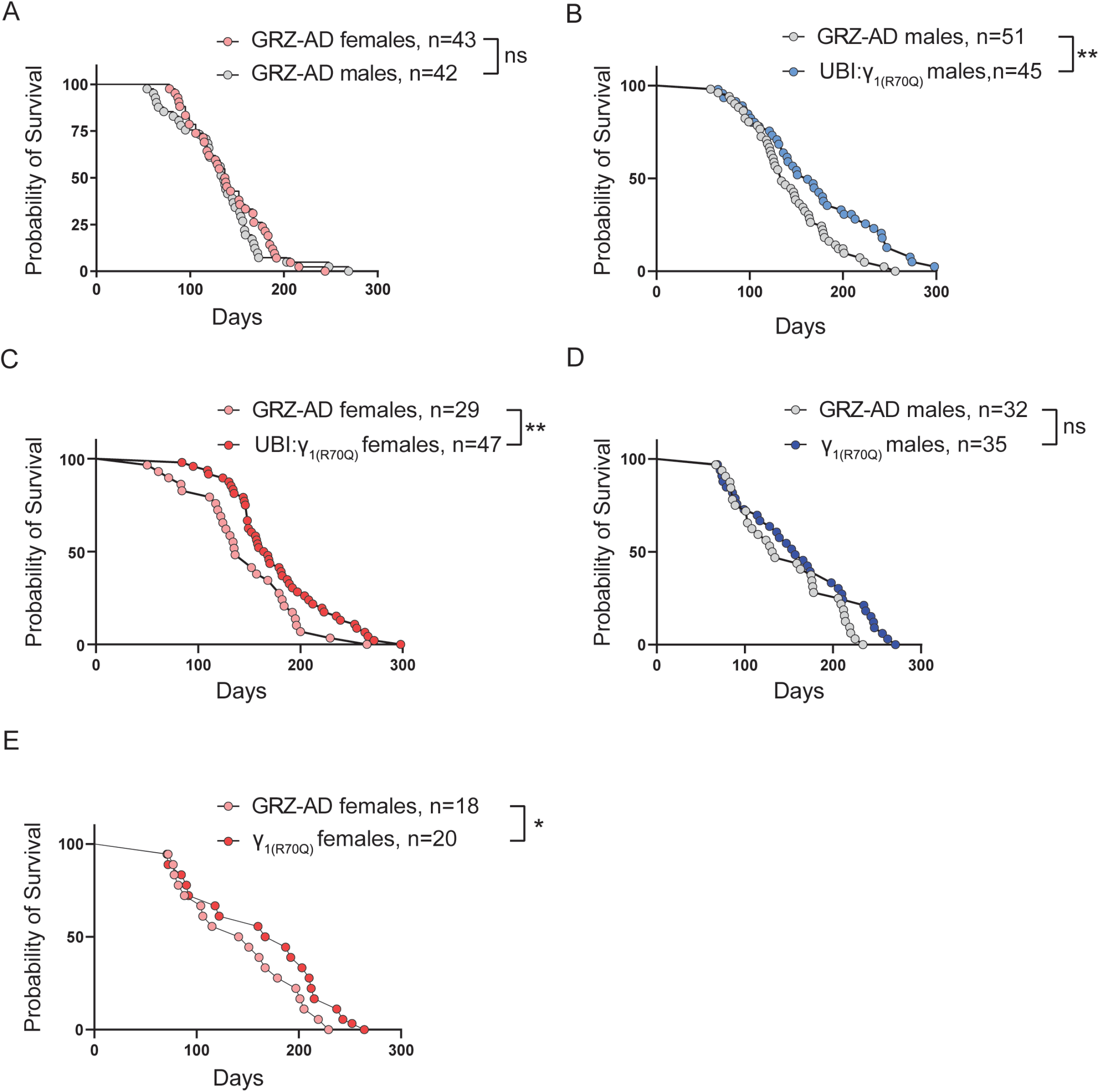
Sustained activation of AMPK_γ1_ promotes longevity. **A**) Survival analysis of wild-type male and female single-housed fish. **B**) Survival analysis of Ubi:γ_1(R70Q)_ compared to wild-type male fish. **C**) Survival analysis of Ubi:γ_1(R70Q)_ compared to wild-type female fish. **D**) Survival analysis of Y_1(R70Q)_ compared to wild-type male fish. **E**) Survival analysis of Y_1(R70Q)_ compared to wild-type female fish. Statistical analysis was calculated by two-tailed log-rank. **\***:P<0,05; **\*\***: P<0,01; **\*\*\***:P<0,001.

**Extended Data Figure 6.**
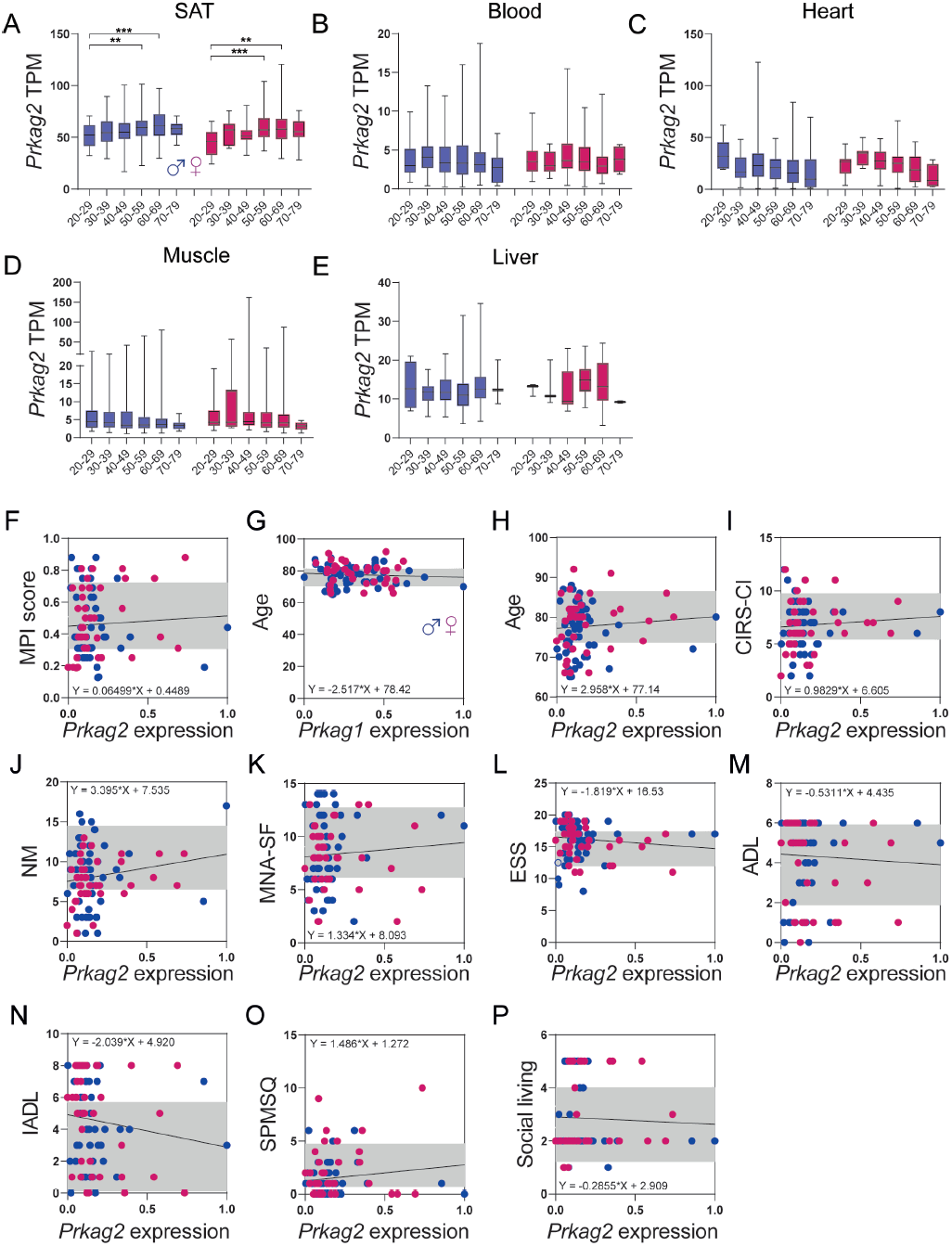
Human AMPK γ1 expression is a biomarker of healthy aging. **A**-**E**) Box plots showing human normalized *Prkag2* expression levels across age groups in decadal brackets. Data are available in the GTEx Consortium (GTEx analysis V8). **F**) Scatter plot showing the relationship between *Prkag2* expression and MPI score in males (blue dots) and females (pink dots). The black line represents a linear model fit with a 95% confidence interval in gray. **G**-**H**) Scatter plot showing the relationship between donor’s age and *Prkag1*(G) and *Prkag2* (H) expression. **I**-**P**) Scatter plots comparing the relationship between *Prkag2* expression and MPI-subdomains. Black lines represent linear model fit with a 95% confidence interval in gray. Significance was obtained by one-way ANOVA followed by Tukey’s post-hoc test in A-E. **\***:P<0.05; **\*\***: P<0.01; **\*\*\***:P<0.001.

**Extended Data Figure 7.**
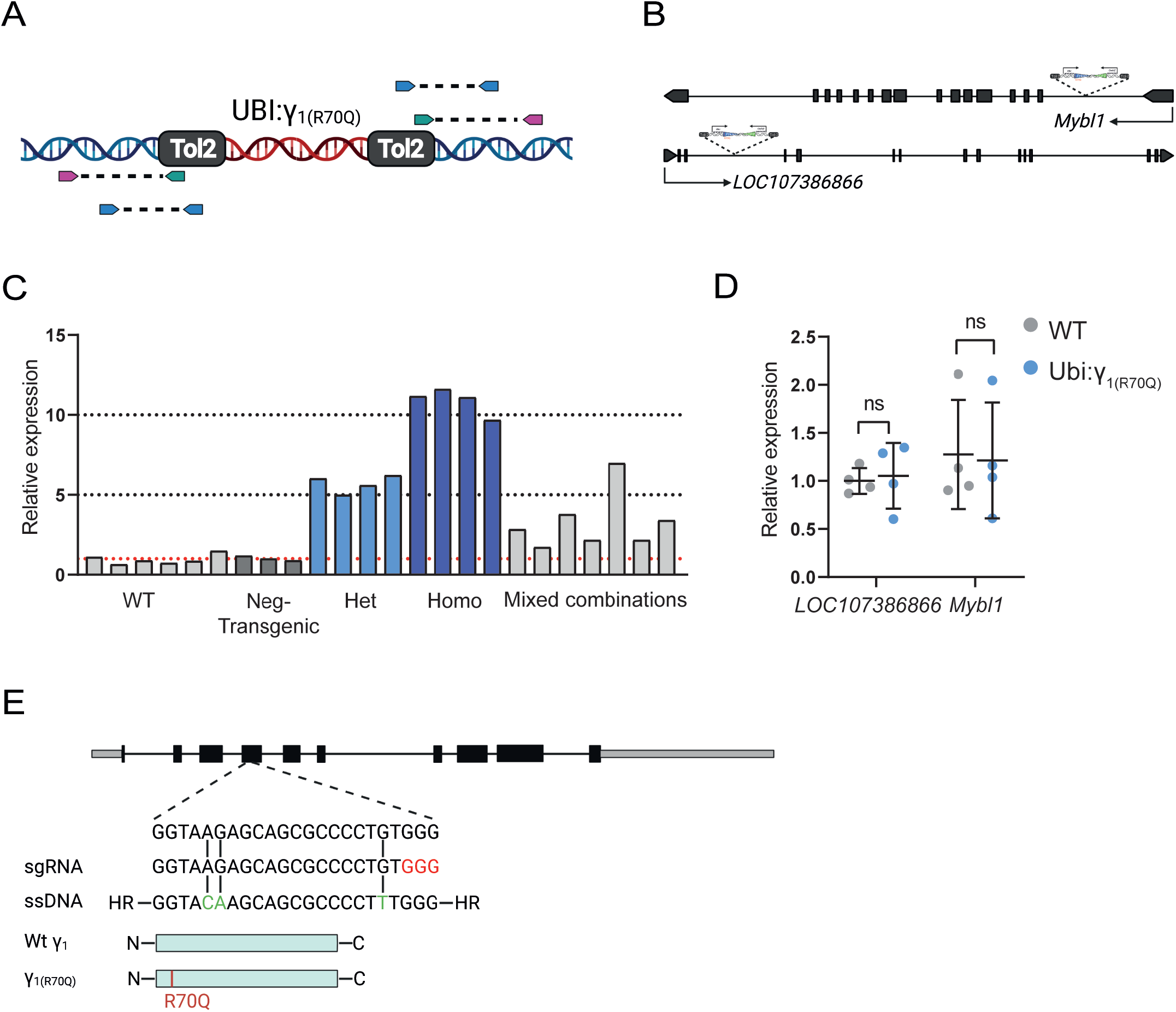
Generation of *Prkag1* mutant lines. **A**) Schematic representation of the transposon integration mapping strategy. The red DNA fragments represent the Ubi:γ_1(R70Q)_ expression cassette, pink arrows represent degenerate primers that anneal to unknown flanking genomic regions. **B**) Representation of the tol-2 expression cassette integration loci. **C**) Fin clip q-PCR analysis of *Prkag1* mRNA expression from fish with different copies of Ubi:γ_1(R70Q)_ expression cassette. Each bar represents an independent biological replicate. **D**) q-PCR analysis of *mybl1* and *LOC107386866* mRNA expression in Ubi:γ_1(R70Q)_ relative to wild-type fish. n=4 fish/group. **E**) Generation of γ_1(R70Q)_ CRISPR mutant line. Location of the sgRNA successfully targeting *Prkag1* exon 4 (Top). Green highlights the precise editing of specific codons introduced by the ssDNA template leading to the R70Q mutation (bottom). Protospace adjacent motif (PAM) is indicated in red. Data in **D**) are presented as mean ± SD. Significance was obtained by nonparametric Mann– Whitney U test.

## Materials and Methods

### Fish husbandry

All experiments were performed on adult (young 6-8 weeks, and old 18-20 weeks old) African turquoise killifish *Nothobranchius furzeri* laboratory strain GRZ-AD. Adult fish were raised singularly in 2.8L tanks from the second week of life. Water parameters were pH: 7-8; kH: 3-5; T: 27°C. The system automatically replaced ten percent of the water with fresh water daily. Fish were raised in 12 hours of light and 12 hours of darkness and fed with 10mg of dry pellet (Biomar inicio plus) and “Premium Artemia Coppens®” twice a day (for a total amount of food daily delivered equal to 2-3% of fish weight). The first feeding was made at 8:30 and the second at 13:30. For tissue collection, fish were sacrificed by rapid chilling. Tissues were rapidly extracted by dissection, snap-frozen in liquid nitrogen, and stored at −80°C. Animal experimentation was approved by “Landesamt für Natur, Umwelt und Verbraucherschutz Nordrhein-Westfalen”: 81-02.04.2019.A055.

### Generation of the Ubi:_γ1(R70Q)_ line

The Tol2 Ubi:H2ACFP-(2x)SV40pA/cmlc2:eGFP vector, originally generated using the TOL-2 kit^48^, was digested with KpnI to excise H2ACFP. Prkag1(R70Q) sequence was generated by PCR-mediated site-directed mutagenesis using the killifish Prkag1 cDNA as a template. Prkag1(R70Q) was then recombined with Ubi:H2ACFP-(2x)SV40pA, previously digested with KpnI to excise H2ACFP, using the NEBuilder® HiFi DNA Assembly Cloning Kit (Primers in supplementary table 13). The resulting plasmid (Ubi:_γ1(R70Q)_) was then amplified in *E. coli* and purified with Wizard® Plus SV Midipreps DNA Purification Promega. Tol2 transposase mRNA was synthesized by in vitro transcription using the mMESSAGE mMACHINE® SP6 (Ambion) kit and the pCS2FA plasmid, previously linearized with NotI, as a template. The mRNA was then purified by lithium chloride precipitation, aliquoted, and stored at -80°C. Transgenic fish were generated by injecting 1-2 nL of a solution containing 30 ng/μl of Tol2 mRNA, 40 ng/μl of Ubi:_γ1(R70Q)_ plasmid, 0.4M of KCl into one cell stage *N. furzeri* embryos and 1% phenol red used as visual control of successful injections. Potential founders expressing a myocardium-specific eGFP signal were identified under a fluorescent microscope and backcrossed with the GRZ-AD wild-type strain for three generations. To map the transposon insertions, we used a PCR-based sequencing approach where forward primers hybridizing at the edges of the tol-2 arms were coupled with degenerate reverse primers to amplify DNA spanning the transposon insertion junctions (Extended Data Figure 7A; supplementary table 13 for primer sequences). All the resulting bands were purified, sequenced, and mapped onto the killifish genome using BLAST. Each integration site was cross-validated using another couple of primers encompassing the transposon insertion junctions (Extended Data Figure 7A). We obtained a stable transgenic line carrying two copies of Ubi:_γ1(R70Q)_: one integrated into the third intron of *mybl* and the other into the first intron of an unidentified gene *LOC107386866*, respectively (Extended Data Figure 7B). We used this dual copy line since a further reduction of copy number resulted in weak transgenic expression in several tissues (data not shown). Fish having two copies of the transgene in heterozygous and homozygous configurations induced overexpression of *Prkag1* of about 5 and 10 folds in the fin (Extended Data Figure 7C). The mRNA expression of resident genes at the integration site appeared unaffected by the transgene insertion (Extended Data Figure 7D). Negative transgenic fish (defined as wild-type in the text) were used as control fish in aging comparative and survival analysis.

### γ1_(R70Q)_ line generation

CRISPR/CAS9 genome editing was performed according to^49^. All the sgRNAs were designed based on CHOP-CHOP web-based tool (https://chopchop.cbu.uib.no). The single-strand DNA (ssDNA) to generate γ1(R70Q) was designed to contain 45bp of homology arm and two base pair mutations in the region encompassing the coding sequence for γ1 amino acid residue R70. Alt-R® S.p. HiFi Cas9 Nuclease, Alt-R® sgRNA, and the ssDNA template were purchased from IDT (see supplementary table 13 for sequences). One-cell stage embryos were injected with 1-2 nl of a solution containing: Cas9 enzyme (200ng/µl), sgRNA (20ng/ul), ssDNA (40ng/ul), KCl (0,2M), 1% phenol red. F0 generation was genotyped by fin-clipping to identify potential founders (Extended Data Figure 7E-7F, supplementary table 13 for primers sequences). Selected founders were then backcrossed with the GRZ-AD strain for four generations to reduce the potential presence of background mutations induced by the CRISPR editing.

### Food consumption analysis

To monitor food consumption, single-housed fish were fed daily with a fixed amount of food (n=40 dry pellets/fish) using a handheld dispensing machine (SDH-1, XQ instruments). Fish were then allowed to eat for one hour. During this time, the water flow was interrupted to prevent food from washing away. At the end of this time, the bottom of every tank was recorded for about 15 seconds by a photo camera (SQ12 spy mini DV camera, RC-group) positioned above the lid. Videos were analyzed to count the number of leftover pellets (LP). Food consumption was calculated as (40-LP)/grams of body weight.

### MicroCT body scan analysis

Euthanized fish were scanned using the high-performance in vivo micro-CT scanner SkyScan 1176 (Bruker) with an isotropic voxel size of 18 µm3 using the following parameters: voltage 45 kV, current 475 µA using a 0.5 mm aluminum filter, and exposure time of 260ms. All the scans were performed over 360 degrees with a rotation step of 0.6 degrees and a frame averaging of 2. Image data were reconstructed using the NRrecon Software (Bruker) with the following parameters: smoothing degree 4, ring artifact reduction 3, beam hardening correction 30%, and defect pixel masking 5%. Grey values of all images were standardized by setting the contrast range of the histogram from 0 to 0.03. Relative sub-cutaneous, visceral fat, and eggs yolk quantification from reconstructed images were determined using the CTAn software (Bruker).

### Tissues fixation, H&E staining, immunofluorescence, EDU labeling, and adipocyte area counting

Killifish were euthanized by rapid chilling and immersed in formalin solution for 72h at 4°C after the visceral cavity was opened, and the gills opercula were removed. Then fish were transferred in EDTA (500mM, Ambion) solution for another 72h and paraffin-embedded. Next, 4µm paraffin sections were made and stored at room temperature. The slides were deparaffinized and stained with Hematoxylin and Eosin using the Gemini station (Thermo Scientific: A81510100). Heat-mediated antigen retrieval was initially used on deparaffinized slides for immunolabellings to break protein cross-links. Next, slides were washed thrice in 1xPBS+triton 0,3%, blocked for one hour at room temperature with a solution of goat normal serum 10% (Abcam: ab748), and BSA 2,5% and incubated with rabbit anti-L-plastin primary antibody (1:400, GTX124420, Genetex) overnight at 4°C. The next day, slides were washed thrice in PBST and incubated first with the SignalStain® Boost IHC Detection Reagent (8114s, CST) for 30 min at room temperature, then with the DAB substrate for 1-2 minutes. Finally, slides were counterstained with hematoxylin, dehydrated with alcohol and xylene, and mounted using Fluoromount-G with DAPI (00-4959-52, Thermo). Images were acquired using Nikon eclipse Ci microscope. To label dividing cells, fish were intraperitoneally injected with 8ug/g of the weight of 5-Ethynyl-2’-Desoxyuridin (EDU) 6 hours before the euthanasia. EDU detection on deparaffinized slides was performed using the Click-iT EDU Alexa Fluor 488 Imaging Kit according to manufacturer instructions (C10337, Thermo). Images were acquired using Leica DMI6000B microscope. The frequency distribution of adipocyte areas was obtained using the Adiposoft software^50^.

### Total RNA extraction and qPCR analysis

Killifish tissues were thawed in RLT buffer (Qiagen) with 1% β-ME v/v and mechanically crushed in a mortar and pestle. They were crushed by plastic beads using Tissue Lyser LT (Qiagen) at 50 oscillations per second for 15 minutes at 4°C. Samples were then allowed to settle for 15 minutes on ice before centrifugation at 16,000g for 10 minutes at 4°C. The supernatant was collected for subsequent RNA extraction using the RNeasy Mini Kit (Qiagen) according to the manufacturer’s instructions. The optional DNAse step was always performed using the RNase-Free DNase Set kit (Qiagen) according to the manufacturer’s instructions. The concentration and purity of the RNA were measured by NanoDrop. cDNA was generated using iScript (Bio-Rad). qRT-PCR was performed with Power SYBR Green (Applied Biosystems) on a ViiA 7 Real-Time PCR System (Applied Biosystems). Four technical replicates were averaged for each sample per primer reaction. EIF3C was used as an internal control for killifish samples, B-actin for Human samples (See supplementary table 13 for primers sequences)

### Transcriptomic profiling and computational analysis

Visceral adipose tissue from fully fed or fasted male fish was collected for expression profiling. To reduce possible batch effects or variability due to circadian rhythms, fish were sacrificed all at once within two hours in the early afternoon. Harvested tissues were snap-frozen in liquid nitrogen and stored at -80°C. RNA extraction of all samples was done at the same time. About 1 μg of total RNA was used per sample for library preparation. The ribosome removal step was done using the RiboZero rRNA removal kit (Illumina). The sequencing was performed on Illumina HiSeq4000 sequencing system (∼30 million reads per sample) using a paired-end two × 100 nt sequencing protocol. After removal of rRNA and tRNAs, reads were pseudo aligned to the reference genome (Nfu_20140520) using Kallisto (0.45.0)^51^. Genes with less than 10 overall reads were removed. After normalization of read counts by making use of the standard median-ratio for estimation of size factors, pair-wise differential gene expression was performed using DESeq2 (1.24.0)^52^. Log2 fold changes were shrunk using approximate posterior estimation for GLM coefficients. The KEGG pathway analysis of significant genes was performed using ShinyGO V0.76.2. FDR value <0,05 was considered to be significant. Hierarchical clustering was calculated using FLASKI (https://flaski.age.mpg.de/). All raw RNA sequencing data can be found in the SRA database, bioProject ID: PRJNA817434.

### Blood glucose, triglycerides, and free fatty acids quantification

Killifish were euthanized by rapid chilling and then transferred on a piece of gauze to carefully dry off their body. A small incision on the lateral side nearby the caudal fin was made using a steel blade. The incision was deep enough to penetrate skin and muscle and reach the aorta to release the blood. Two ul of blood were immediately used to determine blood glucose concentration (mg/dl) using the glucometer Accu-check guide (Roche). In contrast, 8µl were used to determine blood triglyceride concentration (mg/dl) using Accutrend-plus (Roche). The excess blood was collected in a 1.5ml tube. To avoid hemolysis and coagulation, blood was immediately mixed with 0.5 v/v of heparin (3mg/ml in 1XPBS) and kept on ice for 5-10 minutes. Samples were then centrifuged for 10 minutes at 8,000 rpm. Afterward, the plasma was collected, transferred to a new 1.5ml tube, and used directly for analysis or flash frozen in liquid nitrogen and stored at -80°C. 8µl of plasma was used to determine NEFAs concentration using the Free Fatty Acid Quantitation Kit (Sigma: MAK044) according to the manufacturer’s instructions. NEFAs values were expressed as nmol/plasma total protein concentration.

### Western blot analysis

RIPA buffer supplemented with cOmplete Protease Inhibitor (Roche) and PhosSTOP (Roche) was used for total protein extraction. Samples were lysed in RIPA buffer using Bioruptor Sonication System (Diagenode) for 20mins, then kept on ice for another 15 mins. Afterward, samples were centrifuged for 10 min at maximum speed to remove cell debris, and the supernatants were transferred to new tubes. Protein concentration was estimated using micro BCA Protein Assay Kit (Thermo Fisher Scientific). Protein samples were then heated to 75°C for 15 min in 2xLaemmli buffer with 0.9% 2-mercaptoethanol. Next, 20-25ug of protein per sample were loaded on 4–15% MiniPROTEAN TGXTM Precast Protein Gels (Bio-Rad), and electrophoresis was performed at constant voltage of 200V for about 30-40 min. After gel separation, proteins were transferred on nitrocellulose membranes using Trans-Blot TurboTM Transfer System (BioRad) and blocked for 1 hour with 5% Bovine serum albumin (BSA) in (TBST1X). Afterward, the membranes were incubated overnight with primary antibodies and for 1 hour with the secondary antibody. Imaging of the membranes was performed with ChemiDoc Imager (BioRad). Western Lightning Plus Enhanced Chemiluminescence Substrate (PerkinElmer) was used as the chemiluminescence reagent. Here follows the list of antibodies and relative dilution used in this study. Total AMPKα (CST:2532; 1:1000), phosphor (Thr172)-AMPKα (CST:2535; 1:1000), total AMPKβ (CST:4150;1:1000), total-ACC (CST:3676; 1:500), phospho(Ser79)-ACC (CST:3661; 1:1000), γ2 (Invitrogen: PA522331; 1:1000), α-Tubulin (Sigma:T9026, 1:10000).

### Lifespan analysis

All eggs used for survival analysis were collected daily from aged-matched parents within ten days. After hatching, larvae were housed together (4 larvae for 1.1L tank) until three weeks of age, then single-housed in 2.8L tanks for the remaining lifespan. Fish mortality was scored starting in the sixth week when sexual maturation was fully reached. At this point, fish exhibiting reduced size, incomplete sexual maturation, or body malformations were removed from the cohorts. Both males and females were used for the experiments. Fish were examined daily for signs of ill health. Senescent fish (>28 weeks) that showed vital signs of distress such as severe lethargy, anorexia, or advanced sarcopenia were euthanized for humane reasons. The age at which an ill fish was euthanized was taken as the last available estimate of its natural lifespan. Survival curves were calculated using the Kaplan-Meier estimator. Statistical significance was calculated by the Log-rank test.

### Patient population and isolation of human PBMCs

The study recruited participants ≥ 65 years of age who presented to the emergency department at the University Hospital Cologne. Approval was obtained from the institutional review board of the University of Cologne (EK20-1346, EK19-1275), and written informed consent was obtained from all patients. The study was conducted in accordance with the Declaration of Helsinki and the good clinical practice guidelines by the International Conference on Harmonization and was registered in the German Clinical Trials Register (DRKS00024592).

In brief, whole heparinized blood was diluted 1:1 with PBS in sterile conditions and transferred to a Leucosep™ tube (Greiner Bio-One, cat no. GREI163290_500). Following centrifugation, a layer of PBMCs became visible and was carefully aspirated. PBMCs were then counted and assessed for viability using trypan blue staining. Subsequently, PBMCs were cryopreserved in liquid nitrogen.

### RNA extraction from human PBMCS

Aliquots of 1e^6^ viable PBMCs were thawed on ice. RNA was extracted employing a bead-based approach on a KingFisher™ Flex Magnetic Particle Processor (ThermoFisher) using the MagMAX™ mirVana™ Total RNA Isolation Kit (ThermoFisher, cat no. A27828) according to the manufacturer’s instructions. The concentration and purity of the RNA were measured by NanoDrop.

### Multidimensional prognostic index

The Multidimensional prognostic index (MPI) was calculated based on a Comprehensive Geriatric Assessment. The MPI included clinical, cognitive, functional, nutritional, and social parameters and was carried out using six standardized scales (Exton Smith Scale (ESS), Instrumental Activities of Daily Living (IADL), Activities of Daily Living (ADL), Cumulative Illness Rating Scale (CIRS), Mini-Nutritional Assessment Short Form (MNA-SF) and Short Portable Mental Status Questionnaire (SPMSQ), as well as information on the number of medications and social support network, for a total of 63 items in eight domains. An MPI was calculated from CGA as described before^22^, expressing it as a score from 0 to 1, being subdivided into three MPI groups: MPI-1 (robust), 0.0-0.33; MPI-2 (pre-frail), 0.34-0.66; and MPI-3 (frail), 0.67-1.0.

## Notes

### Competing Interest Statement

The authors have declared no competing interest.

### Summary of Updates

Supplementary figure 5 revised Acknowledgments updated

## References

1 Brandhorst, S. et al. A Periodic Diet that Mimics Fasting Promotes Multi-System Regeneration, Enhanced Cognitive Performance, and Healthspan. Cell Metabolism 22, 86–99 (2015). https://doi.org:10.1016/j.cmet.2015.05.012

2 Mitchell, S. J. et al. Daily Fasting Improves Health and Survival in Male Mice Independent of Diet Composition and Calories. Cell Metab 29, 221–228 e223 (2019). https://doi.org:10.1016/j.cmet.2018.08.011

3 Colman, R. J. et al. Caloric restriction delays disease onset and mortality in rhesus monkeys. Science 325, 201–204 (2009). https://doi.org:10.1126/science.1173635

4 Mitchell, S. J. et al. Effects of Sex, Strain, and Energy Intake on Hallmarks of Aging in Mice. Cell Metab 23, 1093–1112 (2016). https://doi.org:10.1016/j.cmet.2016.05.027

5 Hahn, O. et al. A nutritional memory effect counteracts benefits of dietary restriction in old mice. Nat Metab 1, 1059–1073 (2019). https://doi.org:10.1038/s42255-019-0121-0

6 Goodrick, C. L., Ingram, D. K., Reynolds, M. A., Freeman, J. R. & Cider, N. Effects of intermittent feeding upon body weight and lifespan in inbred mice: interaction of genotype and age. Mech Ageing Dev 55, 69–87 (1990). https://doi.org:10.1016/0047-6374(90)90107-q

7 Duriancik, D. M., Tippett, J. J., Morris, J. L., Roman, B. E. & Gardner, E. M. Age, calorie restriction, and age of calorie restriction onset reduce maturation of natural killer cells in C57Bl/6 mice. Nutr Res 55, 81–93 (2018). https://doi.org:10.1016/j.nutres.2018.04.009

8 Morgan E. Levine, J. A. S., Sebastian Brandhorst, Priya Balasubramanian, Chia-Wei Cheng, Federica Madia, Luigi Fontana, Mario G. Mirisola, Jaime Guevara-Aguirre, Junxiang Wan, Giuseppe Passarino, Brian K. Kennedy, Min Wei, Pinchas Cohen, Eileen M. Crimmins,and Valter D. Longo. Low Protein Intake Is Associated with a Major Reduction in IGF-1, Cancer, and Overall Mortality in the 65 and Younger but Not Older Population. Cell Metab (2014). https://doi.org:10.1016/j.cmet.2014.02.006

9 Tonini, C. et al. Effects of Late-Life Caloric Restriction on Age-Related Alterations in the Rat Cortex and Hippocampus. Nutrients 13 (2021). https://doi.org:10.3390/nu13010232

10 Prvulovic, M. R. et al. Late-Onset Calorie Restriction Worsens Cognitive Performances and Increases Frailty Level in Female Wistar Rats. J Gerontol A Biol Sci Med Sci 77, 947–955 (2022). https://doi.org:10.1093/gerona/glab353

11 Hu, C. K. & Brunet, A. The African turquoise killifish: A research organism to study vertebrate aging and diapause. Aging Cell 17, e12757 (2018). https://doi.org:10.1111/acel.12757

12 Harel, I. et al. A platform for rapid exploration of aging and diseases in a naturally short-lived vertebrate. Cell 160, 1013–1026 (2015). https://doi.org:10.1016/j.cell.2015.01.038

13 Cellerino, A., Valenzano, D. R. & Reichard, M. From the bush to the bench: the annual Nothobranchius fishes as a new model system in biology. Biol Rev Camb Philos Soc 91, 511–533 (2016). https://doi.org:10.1111/brv.12183

14 Polacik, M., Blazek, R. & Reichard, M. Laboratory breeding of the short-lived annual killifish Nothobranchius furzeri. Nature Protocols 11, 1396–1413 (2016). https://doi.org:10.1038/nprot.2016.080

15 Miller, K. N. et al. Aging and caloric restriction impact adipose tissue, adiponectin, and circulating lipids. Aging Cell 16, 497–507 (2017). https://doi.org:10.1111/acel.12575

16 Meng, B., Wang, Y. & Li, B. Suppression of PAX6 promotes cell proliferation and inhibits apoptosis in human retinoblastoma cells. Int J Mol Med 34, 399–408 (2014). https://doi.org:10.3892/ijmm.2014.1812

17 Longo, V. D. & Mattson, M. P. Fasting: molecular mechanisms and clinical applications. Cell Metab 19, 181–192 (2014). https://doi.org:10.1016/j.cmet.2013.12.008

18 Lin, J.-R. et al. Rare genetic coding variants associated with human longevity and protection against age-related diseases. Nature Aging 1, 783–794 (2021). https://doi.org:10.1038/s43587-021-00108-5

19 Hamilton, S. R. et al. An activating mutation in the gamma1 subunit of the AMP-activated protein kinase. FEBS Lett 500, 163–168 (2001). https://doi.org:10.1016/s0014-5793(01)02602-3

20 Ou, M. Y., Zhang, H., Tan, P. C., Zhou, S. B. & Li, Q. F. Adipose tissue aging: mechanisms and therapeutic implications. Cell Death Dis 13, 300 (2022). https://doi.org:10.1038/s41419-022-04752-6

21 Defour, M., Michielsen, C., O’Donovan, S. D., Afman, L. A. & Kersten, S. Transcriptomic signature of fasting in human adipose tissue. Physiol Genomics 52, 451–467 (2020). https://doi.org:10.1152/physiolgenomics.00083.2020

22 Pilotto, A. et al. Development and validation of a multidimensional prognostic index for one-year mortality from comprehensive geriatric assessment in hospitalized older patients. Rejuvenation Res 11, 151–161 (2008). https://doi.org:10.1089/rej.2007.0569

23 Schafer, M. et al. Risk Stratification of Patients Undergoing Percutaneous Repair of Mitral and Tricuspid Valves Using a Multidimensional Geriatric Assessment. Circ Cardiovasc Qual Outcomes 14, e007624 (2021). https://doi.org:10.1161/CIRCOUTCOMES.120.007624

24 Veronese, N. et al. Prevalence of multidimensional frailty and pre-frailty in older people in different settings: A systematic review and meta-analysis. Ageing Res Rev 72, 101498 (2021). https://doi.org:10.1016/j.arr.2021.101498

25 Montesano, A. et al. Age-related central regulation of orexin and NPY in the short-lived African killifish Nothobranchius furzeri. J Comp Neurol 527, 1508–1526 (2019). https://doi.org:10.1002/cne.24638

26 Nguyen, H. P. et al. Aging-dependent regulatory cells emerge in subcutaneous fat to inhibit adipogenesis. Dev Cell 56, 1437–1451 e1433 (2021). https://doi.org:10.1016/j.devcel.2021.03.026

27 Ortega Martinez de Victoria, E. et al. Macrophage content in subcutaneous adipose tissue: associations with adiposity, age, inflammatory markers, and whole-body insulin action in healthy Pima Indians. Diabetes 58, 385–393 (2009). https://doi.org:10.2337/db08-0536

28 Bonadonna, R. C., Groop, L. C., Simonson, D. C. & DeFronzo, R. A. Free fatty acid and glucose metabolism in human aging: evidence for operation of the Randle cycle. Am J Physiol 266, E501–509 (1994). https://doi.org:10.1152/ajpendo.1994.266.3.E501

29 Gan, L., Chitturi, S. & Farrell, G. C. Mechanisms and implications of age-related changes in the liver: nonalcoholic Fatty liver disease in the elderly. Curr Gerontol Geriatr Res 2011, 831536 (2011). https://doi.org:10.1155/2011/831536

30 Brook, M. S. et al. Synchronous deficits in cumulative muscle protein synthesis and ribosomal biogenesis underlie age-related anabolic resistance to exercise in humans. J Physiol 594, 7399–7417 (2016). https://doi.org:10.1113/JP272857

31 Kalamakis, G. et al. Quiescence Modulates Stem Cell Maintenance and Regenerative Capacity in the Aging Brain. Cell 176, 1407–1419 e1414 (2019). https://doi.org:10.1016/j.cell.2019.01.040

32 Brunet, A., Goodell, M. A. & Rando, T. A. Ageing and rejuvenation of tissue stem cells and their niches. Nat Rev Mol Cell Bio (2022). https://doi.org:10.1038/s41580-022-00510-w

33 Ocampo, A. et al. In Vivo Amelioration of Age-Associated Hallmarks by Partial Reprogramming. Cell 167, 1719–1733 e1712 (2016). https://doi.org:10.1016/j.cell.2016.11.052

34 Lu, Y. et al. Reprogramming to recover youthful epigenetic information and restore vision. Nature 588, 124–129 (2020). https://doi.org:10.1038/s41586-020-2975-4

35 Hishida, T. et al. In vivo partial cellular reprogramming enhances liver plasticity and regeneration. Cell Rep 39, 110730 (2022). https://doi.org:10.1016/j.celrep.2022.110730

36 Wang, C. et al. In vivo partial reprogramming of myofibers promotes muscle regeneration by remodeling the stem cell niche. Nat Commun 12, 3094 (2021). https://doi.org:10.1038/s41467-021-23353-z

37 Harrison, D. E. et al. Rapamycin fed late in life extends lifespan in genetically heterogeneous mice. Nature 460, 392–395 (2009). https://doi.org:10.1038/nature08221

38 Bitto, A. et al. Transient rapamycin treatment can increase lifespan and healthspan in middle-aged mice. Elife 5 (2016). https://doi.org:ARTNe1635110.7554/eLife.16351

39 Juricic, P. Long-lasting geroprotection from brief rapamycin treatment in early adulthood by persistently increased intestinal autophagy. Nature Aging (2022).

40 Mieulet, V. et al. S6 kinase inactivation impairs growth and translational target phosphorylation in muscle cells maintaining proper regulation of protein turnover. Am J Physiol Cell Physiol 293, C712–722 (2007). https://doi.org:10.1152/ajpcell.00499.2006

41 Selman, C. et al. Ribosomal protein S6 kinase 1 signaling regulates mammalian life span. Science 326, 140–144 (2009). https://doi.org:10.1126/science.1177221

42 Zhang, C. S. et al. Fructose-1,6-bisphosphate and aldolase mediate glucose sensing by AMPK. Nature 548, 112–116 (2017). https://doi.org:10.1038/nature23275

43 Scott, J. W. et al. CBS domains form energy-sensing modules whose binding of adenosine ligands is disrupted by disease mutations. J Clin Invest 113, 274–284 (2004). https://doi.org:10.1172/JCI19874

44 Ross, F. A., Jensen, T. E. & Hardie, D. G. Differential regulation by AMP and ADP of AMPK complexes containing different gamma subunit isoforms. Biochem J 473, 189–199 (2016). https://doi.org:10.1042/BJ20150910

45 Yavari, A. et al. Chronic Activation of gamma2 AMPK Induces Obesity and Reduces beta Cell Function. Cell Metab 23, 821–836 (2016). https://doi.org:10.1016/j.cmet.2016.04.003

46 Kim, M. et al. Mutation in the gamma2-subunit of AMP-activated protein kinase stimulates cardiomyocyte proliferation and hypertrophy independent of glycogen storage. Circ Res 114, 966–975 (2014). https://doi.org:10.1161/CIRCRESAHA.114.302364

47 Astre, G. et al. Sex-specific regulation of metabolic health and vertebrate lifespan by AMP biosynthesis. bioRxiv, 2022.2001.2010.475524 (2022). https://doi.org:10.1101/2022.01.10.475524

48 Kwan, K. M. et al. The Tol2kit: a multisite gateway-based construction kit for Tol2 transposon transgenesis constructs. Dev Dyn 236, 3088–3099 (2007). https://doi.org:10.1002/dvdy.21343

49 Harel, I., Valenzano, D. R. & Brunet, A. Efficient genome engineering approaches for the short-lived African turquoise killifish. Nat Protoc 11, 2010–2028 (2016). https://doi.org:10.1038/nprot.2016.103

50 Galarraga, M. et al. Adiposoft: automated software for the analysis of white adipose tissue cellularity in histological sections. J Lipid Res 53, 2791–2796 (2012). https://doi.org:10.1194/jlr.D023788

51 Bray, N. L., Pimentel, H., Melsted, P. & Pachter, L. Near-optimal probabilistic RNA-seq quantification. Nat Biotechnol 34, 525–527 (2016). https://doi.org:10.1038/nbt.3519

52 Love, M. I., Huber, W. & Anders, S. Moderated estimation of fold change and dispersion for RNA-seq data with DESeq2. Genome Biol 15, 550 (2014). https://doi.org:10.1186/s13059-014-0550-8

